# How Demographic Noise Shapes Phenotypic Clusters in Environmental Gradients

**DOI:** 10.64898/2026.05.14.725167

**Authors:** Nathanaël Boutillon, Louise Fouqueau

## Abstract

2

Although resources are typically distributed continuously in space, the distribution of species is often organized in discrete clusters. Such clusters have been shown to spontaneously arise in population densities, even in environments with continuously varying conditions, under the phenomenon known as Turing instability. In this work, we consider two models grounded in population dynamics: a one-dimensional model based on the nonlocal Fisher-KPP equation, and a two-dimensional model involving an environmental gradient. We show that phenotypic clusters emerge in these models, and we prove that they do not emerge because of Turing instability, but because of stochasticity. We first consider initial populations that are uniformly distributed in the state space. We show that phenotypic clusters quickly emerge, and that the distances between them depend on population size, that is, on the degree of stochasticity. In addition to the effect of population size, we provide quantitative estimates of the various parameters of the model with an environmental gradient on the equilibrium distance between phenotypic clusters. Next, we start from clearly defined phenotypic clusters and modify the distance between those. We identify three regimes in the connection between population size, the initial distances between clusters, and the distances between clusters at equilibrium. Last, on the two-dimensional model, we relax the hypothesis of complete clonality by varying the effective recombination rate. We explore its effect on phenotypic clustering, and show that phenotypic clustering decays drastically with slight recombination.

**Non-specialist summary:** We explored the origin of discrete phenotypic clusters in populations living within a continuous space. Unlike previous studies, our research highlights the importance of stochasticity in the emergence of discreteness. We demonstrate this in two types of model: the first considers a single dimension corresponding either to the phenotype or to the resource spectrum, while the second considers a geographic dimension and a phenotypic dimension. We demonstrate how population size and the other model parameters affect the stable minimum distance between coexisting clusters, as well as how the initial distance between the phenotypes influences this distance. Finally, we relax the hypothesis of complete clonality by introducing sexual reproduction and varying an effective recombination rate.

## 3 Introduction

The concept of *competitive exclusion* refers to the principle that two species cannot stably coexist when they occupy overlapping ecological niches and compete for limited resources. Given this principle, some studies in theoretical ecology aimed at understanding’species packing’, that is, how densely species can be ‘packed’ in a one-dimensional phenotypic space (May 1969). In this framework, May & MacArthur (1972) proved that in a constant and uniform environment (i.e. when the quantity of resource abundance does not change over time and space), there is paradoxically no limit in the packing density. On the other hand, if resource abundance varies stochastically in time, then packing density is strongly limited, even for vanishingly small noise.

In mathematical models, including the model of May & MacArthur (1972), overlapping ecological niches are encoded via *non-local competition*. A systematic mathematical study of this phenomenon was initiated in a work by Britton (1989), where a non-local competition kernel was incorporated into a standard population growth model with mutation. Subsequent studies on the model introduced by Britton, henceforth called ‘phenotype-only model’, as well as some variants, have demonstrated that some shapes of the competition kernel lead to *Turing patterns* that take the form of clusters in the phenotypic space (e.g., Sasaki 1997; Genieys *et al*. 2006; Berestycki *et al*. 2009; Hernández-García *et al*. 2009). Independently, Rogers *et al*. (2012) identified demographic noise as an additional mechanism driving phenotypic clustering, including in situations where the competition kernel does not produce Turing patterns.

The phenomenon of phenotypic clustering has also been explored in population models with an additional structure in geographical space (Doebeli & Dieckmann 2003; Leimar *et al*. 2008). In these models, each geographic location is associated with an optimal phenotype that changes linearly with the position. This linear dependency is known as an environmental gradient and corresponds to a framework commonly used in population genetics to study, for instance, the emergence of genetic clines (Haldane 1948; Barton 1999), range limits and range expansion (Kirkpatrick & Barton 1997; Polechová & Barton 2015), or geographic parthenogenesis (Fouqueau & Roze 2021). Using an adaptive dynamic framework, Doebeli & Dieckmann (2003) explored how the steepness of the gradient and the distance of migration in geographical space influence evolutionary branching, which in the long run leads to distinct phenotypic clusters. The mechanism behind this clustering has remained contested: it was first attributed to edge effects (Polechová & Barton 2005), but this explanation was disputed by Leimar *et al*. (2008), which attributed clustering to Turing instability. However, as we will demonstrate, no Turing instability can occur in the situation studied in Doebeli & Dieckmann (2003). In fact, these explanations fall short because they are formulated within deterministic frameworks; yet stochasticity had already been identified as a strong constraint on species packing (May & MacArthur 1972). Adapting this idea to the gradient setting, we will show that the clustering observed in Doebeli & Dieckmann (2003) is in fact a consequence of demographic noise.

Overall, the role of stochasticity in the emergence of phenotypic clusters has been established, as far as we know, only in the phenotype-only model (Rogers *et al*. 2012). Our goal in this work is to expand the study initiated in Rogers *et al*. (2012), and to extend it to the gradient model. We will first focus on the phenotype-only model, and we will show that phenotypic clustering can indeed arise without Turing instability. Through analytical studies and simulations, we will prove that phenotypic clustering is a consequence of stochasticity, and that the packing density depends on population size, thereby illustrating the effect of stochasticity.

Secondly, we will consider a specific form of gradient model of a population structured along one-dimensional geographical and phenotypical spaces. We will consider a stabilizing selection towards an optimal phenotype that changes linearly with the geographical position. We will also include density-dependence with a constant carrying capacity across space. Using simulations, we will see that for sufficiently low population densities, phenotypic clusters also emerge in the gradient model. We will show that the gradient model is in fact closely related to the phenotype-only model, thereby allowing us to extrapolate the analytical results obtained for the phenotype-only model to the gradient model.

Finally, we will relax the assumption of complete clonality by introducing sexual reproduction in the gradient model. We will explore how the effective rate of recombination, which is defined as the product of the rates of sex and recombination, changes the results of phenotypic clustering. Our results show that clustering disappears quickly even with a slight increase in the effective recombination rate.

## 4 Methods

**Table 1.**
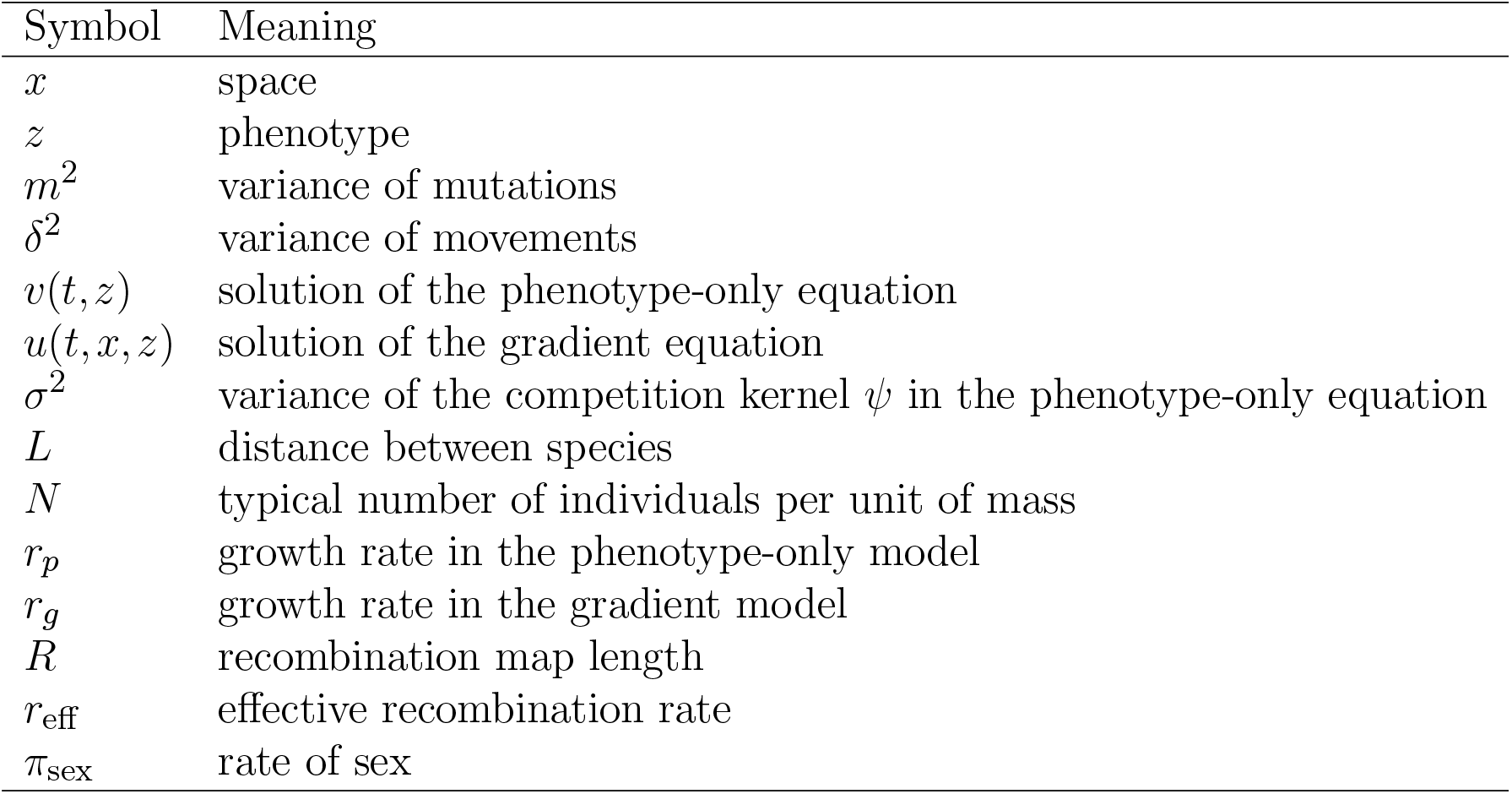
Overview of used notations.

### 4.1 Description of the phenotype-only model

#### Deterministic model

First, we consider a clonal population in which each individual is characterized by a phenotype belonging to ℝ. The population is characterized by the density *v*(*t, z*) of individuals carrying the phenotype *z* ∈ ℝ, which evolves over a continuous time *t* ∈ [0, +∞).

We assume that individuals have an intrinsic growth rate *r*_*p*_ > 0, and are submitted to competition between each other. The strength of competition between two individuals of types *z* and *z*′ is considered to be proportional to *ψ*(*z* −*z*′), for some function *ψ* called the *competition kernel*. We assume that *ψ* is a positive function, that *ψ*(*z*) decreases as |*z*| increases, that *ψ*(*z*) = *ψ*(−*z*), and that 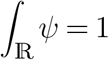. In other words, competition becomes stronger as the phenotypes get closer. Clonality implies that the only difference between the phenotypes of the parent and the offspring is due to *mutations*. We suppose that mutations follow a diffusion with variance *m*^2^: reproduction occurs on a short timescale, and has a weak effect proportional to *m*.

The population is described by the following reaction-diffusion equation:

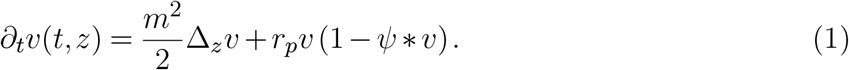

Here, *ψ* * *v* is the convolution product between *ψ* and *v*(*t, ·*), defined by

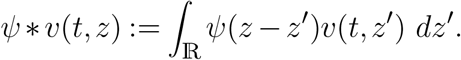

The convolution product translates the fact that individuals with phenotype *z* suffer from the competition pressure induced by individuals with phenotype *z*′ with a strength *ψ*(*z* − *z*′). To obtain Equation (1), we renormalized the population in such a way that the carrying capacity (i.e., the total mass that the environment is able to carry) is equal to 1.

#### Stochastic simulations

Equations similar to (1), and stochastic variants, have been derived from individual-based models in Champagnat *et al*. (2006) (see in particular their Section 2 and Theorem 4.5); we performed stochastic simulations based on those individual-based models.

In order to perform simulations, some approximations were taken. First, we considered discrete time in the simulations while (1) assumes continuous time. Second, (1) assumes continuous phenotype in the unbounded space ℝ. In our simulations, however, we restricted the unbounded space ℝ to a bounded interval [0, *n*_*p*_ − 1], and discretized the phenotypes into a finite number *n*_*p*_ of admissible phenotypes. In other words, each phenotype *z* belongs to {0,…, *n*_*p*_ − 1}, *n*_*p*_ ≫ 1. We point out that considering finitely many phenotypes allows us to handle extremely large populations, by considering the number of individuals within each phenotype, instead of considering each individual separately.

A difficulty with the restriction of the phenotypic space is that the boundary may have an important effect on the dynamics: individuals near the boundaries would be submitted to a much weaker competition than those having an intermediate phenotype, and would thus have a fitness advantage. As we wish to focus on the effects of stochasticity, it is important to avoid boundary effects. We achieved this by considering a periodic phenotype: namely, phenotype *z* = *z*′ + *kn*_*p*_ is identified with phenotype *z*′ for each *k* ∈ ℤ. In this periodic setting, the distance between two phenotypes *z, z*′ ∈ {0,…, *n*_*p*_ − 1} (which determine the strength of the competition between *z* and *z*′) is

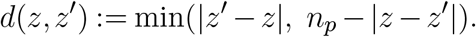

For each *z* ∈ {0,…, *n*_*p*_ − 1}, let *n*_*z*_ be the number of individuals carrying phenotype *z*. We simulate *n*_*z*_, which approximates the population size *Nv*_*z*_. The simulation starts from *n*_*z*_ ≡ *N* individuals at each phenotype *z*, for some ‘typical population size’ *N* . Then, at each time step (of length *dt*) of the simulation, the following takes place:

1. Each individual gives birth to an offspring with probability *dt* × *r*_*p*_. This corresponds to the growth term *r*_*p*_*v* in (1).
2. An individual with phenotype *z* living in deme *x* ∈ {1,…, *n*_*p*_} dies with probability 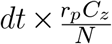, where

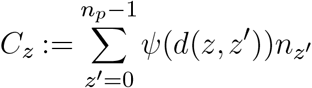

is the competition suffered by this individual. This corresponds to the competition term −*r*_*p*_*ψ* \**v* in (1).
3. The phenotype *z*′ of the offspring depends on the phenotype *z* of its parent: *z*′ equals *z* − 1 or *z* + 1 with equal probability *dt* × *m*^2^/2, and *z*′ = *z* with probability 1 − *dt* × *m*^2^. This corresponds to the diffusion term 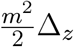 in (1).

#### Gaussian approximation

For the subsequent mathematical analysis, we formally approximated our stochastic simulations with the following equation:

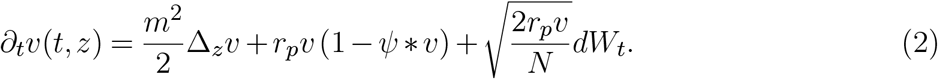

In (2), *dW*_*t*_ is space-time white noise, and *Nv* is to be understood as a number of individuals.

We performed simulations with both approaches: the mechanistic one described above, and with the Gaussian approximation (2). They yielded no qualitative differences. To understand how (2) approximates our stochastic simulations, we note that the number of births in our stochastic simulations is a binomial law *Binom*(*Nv*_*z*_, *dt r*_*p*_), and that the number of deaths is 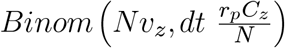. By the central limit theorem, therefore, the net population growth is approximately (for small *dt* and large *N*) a Gaussian law with mean 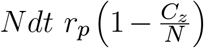 and variance 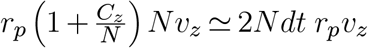. The mean of the Gaussian is encoded by the *r*_*p*_*v* (1 −*ψ* \**v*) in (2). The variance is encoded by the 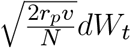 term in (2).

We emphasize that the noise term in (2) is different from the noise term usually used in population genetics, which is of the form 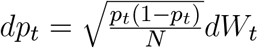. In population genetics, *p*_*t*_ is an allele proportion within a population, whereas here *v*_*t*_ is the size of the entire population. To understand this distinction, let 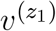 and 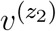 be two independent subpopulations (with different phenotypic values *z*_1_ and *z*_2_) that solve 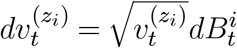, for two independent white noises *dB*^*i*^, *i* = 1, 2. Let *v* := *v*^1^ +*v*^2^ be the total population size and 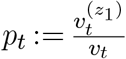 be the proportion of type Praba 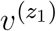 . Then one has 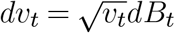 and 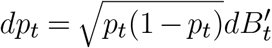 for some well-chosen white noises *dB* and *dB*′.

#### Mean spacing

We estimated the mean spacing between successive phenotypes by identifying low-density regions below a threshold (set to one-tenth of the maximum density), counting the number of such gaps after filtering out small ones, and dividing the domain length by the number of gaps.

### 4.2 Description of the gradient model

#### Deterministic model

Second, we consider a population of individuals living in a linear environment, where each position is associated with an optimal phenotype. Each individual carries a phenotype *z* ∈ ℝ which is submitted to stabilizing selection of strength 1/*V*_*s*_. The optimal value of the phenotype changes linearly with slope *b* > 0; in other words, in position *x* ∈ ℝ, the most adapted phenotype is *z* = *bx*. The maximum fitness of an individual is *r*_*g*_ > 0, and is reduced by maladaptation and competition. Maladaptation is proportional to (*z* −*bx*)^2^/2*V*_*s*_, i.e., a deviation *z* from the optimal phenotype *bx* has a cost *z*^2^/2*V*_*s*_. Competition depends on the total number of individuals in position *x*, called *ρ*(*t, x*). Individuals move in space according to a diffusion with variance *δ*^2^ and mutate according to a diffusion with variance *m*^2^.

As in the phenotype-only model, the population is characterized by the density *u*(*t, x, z*) of individuals carrying phenotype *z* at position *x*. The dynamics of *u* takes place in continuous time, and is described by the following reaction-diffusion equation:

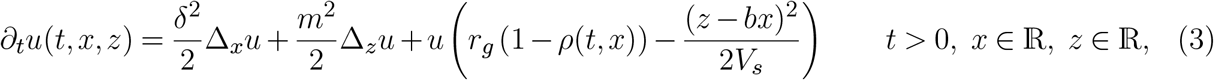

where

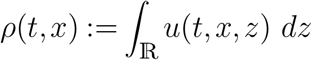

is the total population size at position *x*. We again normalize the population in such a way that the carrying capacity is equal to 1.

The gradient equation (3) and its variants have received some interest in mathematical works (e.g., Champagnat & Méléard 2007; Alfaro *et al*. 2013, 2017; Boutillon 2025). In particular, Alfaro *et al*. (2013) give a criterion based on the principal eigenvalue of an elliptic operator to determine whether the population persists (by invading the whole space) or goes extinct. In our case, the criterion reads:

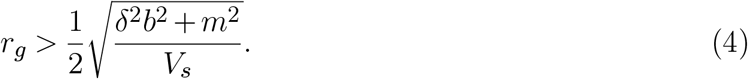

Criterion (4) means that for a population to persist, the intrinsic growth rate *r*_*g*_ must be sufficiently large with respect to the slope of the gradient *b*, the variance of movements *δ*^2^ and the strength of selection 1/*V*_*s*_ (a similar expression appears in Kirkpatrick & Barton 1997 in the case of sexual reproduction. The authors show that the transition between infinite adaptation, limited adaptation, and population extinction depends on the compound parameter 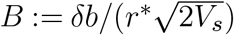, with *r*^*^ corresponding approximately to the growth rate *r*_*g*_. In case of low mutational variance, i.e., *m* ≃ 0, our persistence criterion then reads *B* < 1).

#### Stochastic simulations

In order to perform simulations, we had to circumvent the same difficulties as for simulations of (1). First, (5) assumes continuous time, which we discretized (again with a timestep *dt* ≪ 1). Second, (5) is set in an unbounded environment ℝ with continuous phenotype ℝ. We restricted the unbounded space ℝ and the unbounded phenotype space ℝ into bounded intervals, [0, *n*_*d*_ − 1] and [0, *n*_*p*_ − 1] respectively, and discretized both intervals into a finite numbers *n*_*d*_ of demes and a finite number *n*_*p*_ of admissible phenotypes. By doing so, each position *x* now belongs to a deme in {0,…, *n*_*d*_ − 1} (with *n*_*d*_ ≫ 1), and each phenotype *z* belongs to {0,…, *n*_*p*_ − 1} (with *n*_*p*_ ≫ 1).

Again, a difficulty with such restrictions to bounded spaces of the phenotypic space is that the boundary may have an important effect on the dynamics. In order to avoid these boundary effects, we considered a periodic environment with a periodic phenotype. In order to ensure invariance by translation of our system, we need to make additional approximations. We considered a width *w*, with 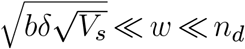, which is understood as the maximal width of the population range around the environmental gradient. We then identify each position/phenotype coordinate (*x, z*) ∈ ℤ^2^ with a position/phenotype coordinate (*x*′, *z*′) such that 0 ⩽ *x*′ ⩽ *n*_*d*_ −*w*− 1 and 0 ⩽ *z*′ ⩽ *n*_*p*_ − *bw* − 1. This identification is illustrated in Figure 1. More formally, we obtain *x*′ by taking the remainder in the Euclidean division of *x* by *n*_*d*_ − *w*, and we similarly obtain *z*′ by taking the remainder of *z* by *n*_*p*_ −*bw*. Such an overlap is necessary to ensure that all individuals (in particular those near the bottom-left and top-right corners of the rectangle) have access to a range of phenotypes around the optimal one with width at least *w*. Since 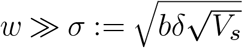, where *σ*^2^ is the typical variance of the population distribution around the optimal phenotype (see Section 5.2), all individuals have access to a similar range of phenotypes around the optimal one, whatever their position in the rectangle is.

**Figure 1.**
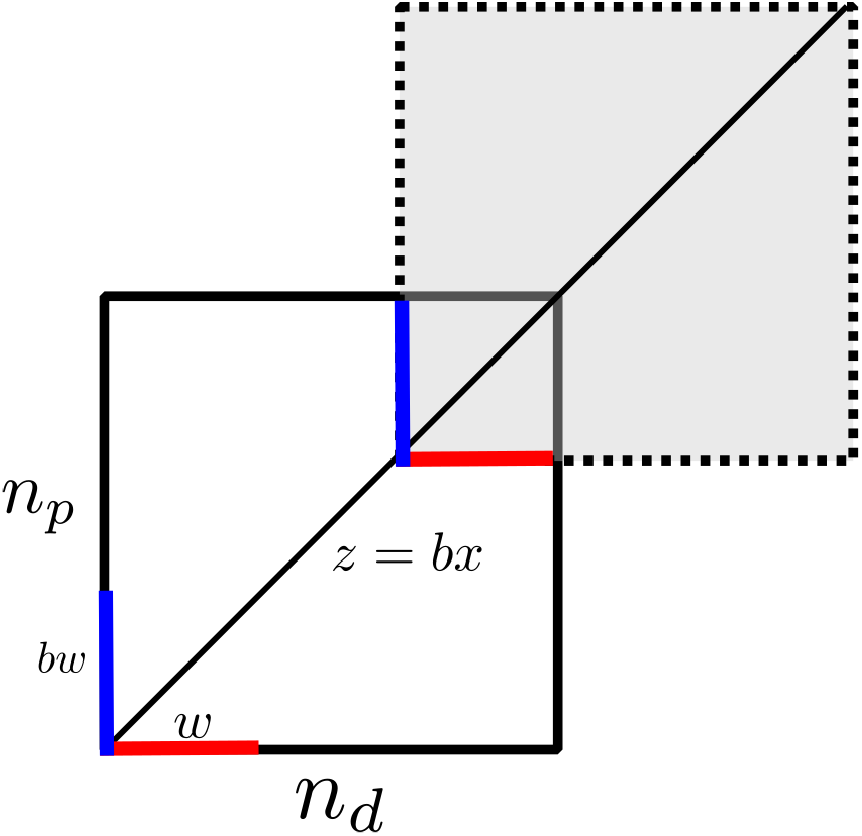
Gradient model in the periodic setting. The horizontal direction is the geographical space, the vertical direction is the phenotypical space. Each individual is characterized by an element of the white area of the bottom-left rectangle. After a mutation or a movement, an individual could leave this area. If it crosses a thick blue line, it jumps instantly to the other blue line and goes on its way. Similarly for the thick red lines. If an individual tries to leave the rectangle by its black boundary, it dies. The width *w* is chosen to be sufficiently large so that very few individuals can reach the black boundary of the rectangle.

For each *x* ∈ {0,…, *n*_*d*_ − 1} and each *z* ∈ {0,…, *n*_*p*_ − 1}, we let *n*_(*x,z*)_ be the number of individuals at position *x* carrying the phenotype *z*. We simulate *n*_(*x,z*)_, which approximates the population size *Nu*_(*x,z*)_. The simulation starts from *n*_(*x,z*)_ ≡ *N* individuals at each position *x* and phenotype *z*. Then, at each time step (of length *dt*) of the simulation, the following takes place:

1. Each individual gives birth to an offspring with probability *dt* × *r*_*g*_. This corresponds to the growth term *r*_*g*_*u*.
2. An individual with phenotype *z* ∈ ℝ living in deme *x* ∈ {0,…, *n*_*d*_ − 1} dies with probability

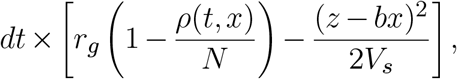

where

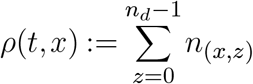

is the total population living in deme *x* at time *t*. This corresponds to the competition and fitness terms 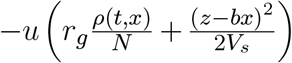.
3. The phenotype *z*′ of the offspring depends on the phenotype *z* of its parent: *z*′ equals either *z* − 1 or *z* + 1 with equal probability *dt* × *m*^2^/2, and *z*′ = *z* with probability 1 − *dt* × *m*^2^. This corresponds to the diffusion term 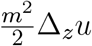. Since the phenotype is periodic, we identify *z*′ with its projection in {1,…, *n*_*p*_} and we change the phenotype according to the method described above.
4. Each offspring of each parent moves to a deme *x*′ depending on the deme *x* of its parent: *x*′ = *x*− 1 with probability *dt* ×*δ*^2^/2, *x*′ = *x* + 1 with probability *dt* ×*δ*^2^/2, and *x*′ = *x* with probability 1 − *dt* ×*δ*^2^. This corresponds to the diffusion term 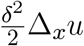. Since space is periodic, we identify *x*′ with its projection in {1,…, *n*_*d*_} and we change the phenotype according to the method described above.

#### Gaussian approximation

As in the phenotype-only model, our stochastic simulations can be formally approximated by a version of (3) with a white-noise term:

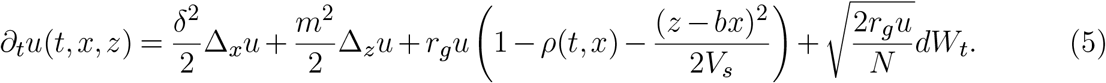

Note that the correspondence between our simulations and (5) is only formal, as solutions of (5) are not well-defined.

Again, *dW*_*t*_ is a white noise in (*x, z*), and *N* ≫ 1 is a scaling parameter corresponding to the typical amount of individuals in one unit of mass. The noise term 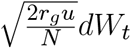 can be understood as in the phenotype-only model above.

#### About periodicity

Note that our simulations are periodic, but the equations (1)–(5), as they are written, are not periodic. Let us explain why this makes little difference. The deterministic equations (1) and (3) have unique solutions (see Prevost 2004; Boutillon & Rossi 2025 for variants of Equation (3)). Hence, the invariance by translation of these equations implies that if the initial condition is periodic, then the solution remains periodic. Therefore, it is almost equivalent to work in the periodic or in the non-periodic setting. Keeping this in mind, we will later work in whichever setting is more convenient.

### 4.3 Sexual reproduction in the gradient model

Lastly, we consider the effect of sexual reproduction in the gradient model. To this aim, we need to clarify the genetic architecture of the quantitative trait submitted to selection. The set-up is equivalent to the one in Fouqueau & Roze (2021). We summarize here the set-up of the individual-based simulation but we refer the readers to the methods section of Fouqueau & Roze (2021) to get the full details. The only difference from this latter work is that the carrying capacity was maintained constant throughout the space (as we do not want to impose any range limits).

We consider haploid hermaphrodite individuals, and suppose that the trait value is the sum of *L* diallelic genes, where the two alleles have allelic effects 0 and *α*, in such a way that the effect of stabilizing selection acting at the gene-level is given by *s* = *α*^2^/2*V*_*s*_. We assume that the *L* loci are evenly spaced along the chromosome, and we ensure that there are sufficiently many loci for some individuals to be perfectly adapted everywhere along the gradient (i.e., from 0 to *b* × *n*_*d*_/*α*). We start from a situation where individuals are well adapted to their positions by setting up genetic clines (Haldane 1948). In addition to the genes coding for the quantitative trait, we consider a modifier locus for the rate of sex. An infinite number of alleles could segregate at this locus (as in a continuum-of-allele model), ranging from 0 (completely clonal) to 1 (completely sexual). This modifier locus affects the investment of individuals (noted *π*_sex_) in each of the two reproductive systems and is held constant throughout the simulation. When an individual reproduces sexually, the other parent is picked from the same deme with a probability proportional to its relative fitness. Self-fertilization is possible. Fertilization is immediately followed by meiosis to produce haploid juveniles and recombination occurs according to map length *R*, corresponding to the average number of cross-overs at meiosis.

We explored the effect of the effective recombination rate *r*_eff_, defined as the product between the rate of sex (*π*_sex_) and the rate of recombination (Roze 2014). The rate of recombination was measured between two genes separated by 100bp and was obtained from the map length *R* using Haldane’s mapping function (Haldane 1919). The values of the map length that were used are: 0.1, 1, 10, 100 and 1000, leading to *r*_eff_ ranging from 4 × 10^−6^ to 0.5. All simulations were run for 50,000 generations.

In this new setting, clusters are not as pronounced as in the asexual setting; therefore, we could not use the same methods to identify spacing. Instead, we considered the phenotypic variance (*V*_*z*_) and the deviation of the mean phenotype from the local phenotypic optima (E[*z*] − *bx*, where the average is taken within deme *x* and will be written 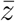 from now on) as in Fouqueau & Roze (2021). To quantify spatial fluctuations of *V*_*z*_ and 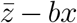, we analyzed the signal using the Fourier transform to decompose it into spatial oscillation modes. The dominant mode, corresponding to the strongest periodic component, was used to estimate the characteristic spatial period as the inverse of its frequency. The main period and amplitude of the fluctuations were obtained from the strength of the dominant Fourier component, which reflects the typical magnitude of the oscillations in space. This method enables us to capture the scale and intensity of spatial structure in a straightforward and reliable manner.

Ten replicates were run for each set of recombination rate and map length values, from which we measured the average and standard error of the period and the amplitude of the genetic variance and the deviation of the local phenotypic optima, as well as the size of the steps.

## 5 Results

### 5.1 Evenly-spaced patterns emerge for small populations

Figures 2–4 show the outcomes of our simulations. In the phenotype-only model, we used a Gaussian competition kernel *ψ*. We observe that when the population size *N* is small (*N* = 20 in the simulation), evenly-spaced peaks emerge in the population density *v*(*t, z*) (Figure 2, left). Those peaks are understood as ‘phenotypic clusters’. Figure 3 shows that the phenotypic clusters arise very quickly in time when started from a uniformly distributed population density. On the other hand, when *N* is large (*N* = 10^10^ on the simulations), no phenotypic clusters arise (Figure 2, right).

**Figure 2.**
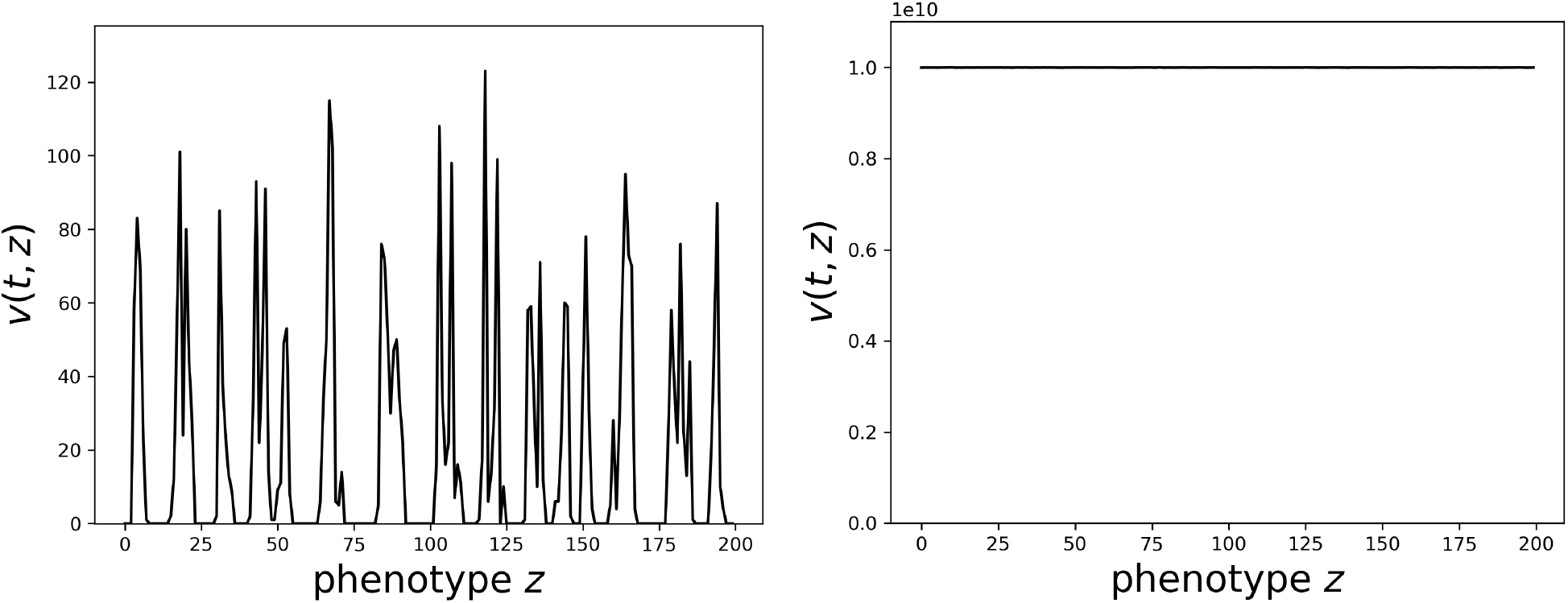
Simulations of the phenotype-only model. In the **left** figure: small population (*N* = 20) corresponding to a stochastic dynamics; in the **right** figure: large population (*N* = 10^10^) corresponding to an almost deterministic dynamics. In both figures, the vertical axis corresponds to the population size *Nv* of each phenotype *z* at time *t* = 1000. The competition kernel *ψ* is a Gaussian with variance *σ*^2^ := 100. The values of the other parameters are: *dx* = 1, *dt* = 0.1, *r*_*p*_ = 1, *n*_*d*_ = 200, *m* ≃ 0.14 (see attached Notebook for exact values). At the beginning of the simulations, the population is uniformly distributed.

**Figure 3.**
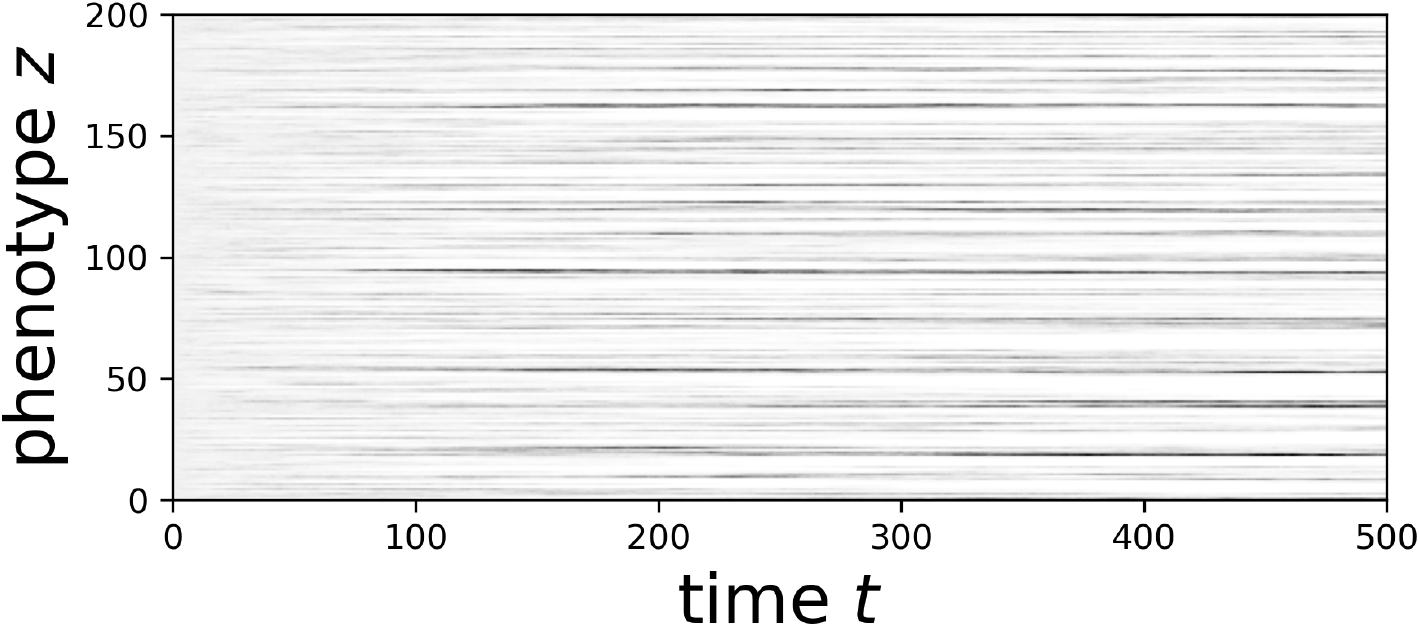
Simulations of the phenotype-only model. Dark shades show high-density phenotypes. The left part of the figure (small times) is homogeneous, but clear patterns quickly emerge. The parameters are the same as in Figure 2, left. A similar figure appears in Rogers *et al*. (2012).

**Figure 4.**
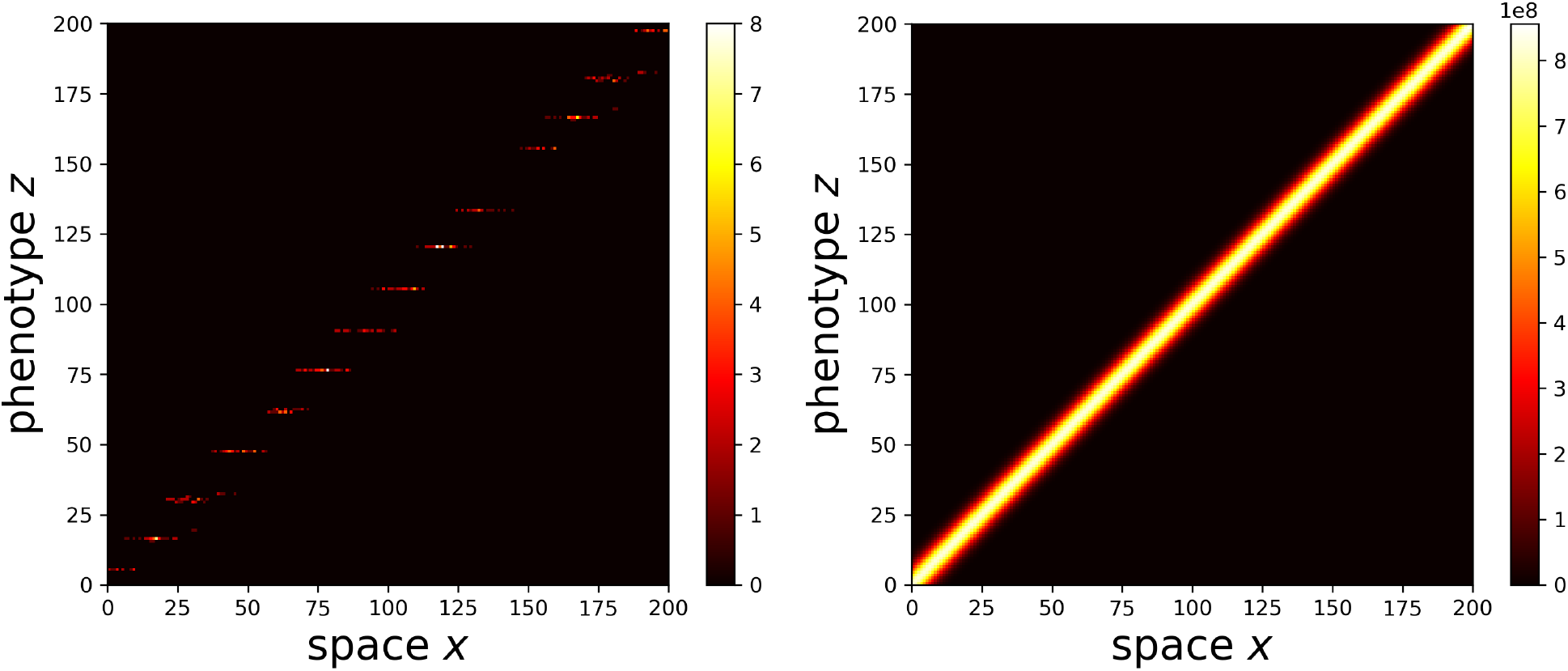
Simulations of the gradient model. The horizontal axis correspond to the geographical space *x*, and the vertical axis to the phenotypic space *z*. As in Figure 2, the **left** figure corresponds to a small population size (*N* = 4) and the **right** figure corresponds to a large population size (*N* = 10^10^). In both figures, we illustrate the density of individuals having phenotype *z* in position *x* at the end of the simulation (time *t* = 1000*N*). The values of the other parameters are: *δ* ≃ 3.16, *m* ≃ 0.14, *V*_*s*_ = 62.5, *b* = 1, *dx* = 1, *dz* = 1, *dt* = 0.1, *r*_*g*_ = 1, *n*_*p*_ = *n*_*d*_ = 200, in such a way that 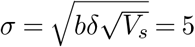 (see attached Notebook for exact values). At the beginning of the simulation, the population is distributed uniformly along the gradient.

Figure 4 shows the result for the gradient model. As in the phenotype-only model, we observe that when population size *N* is small (*N* = 4 in the simulation), ‘stair steps’ patterns arise, corresponding to phenotypic clusters. On the other hand, when *N* is large (*N* = 10^10^ in the simulations), there are no such phenotypic clusters.

These observations suggest that in both models, the emergence of clusters is based on the effect of stochasticity. More precisely, they suggest that clusters appear only at timescales on which stochasticity plays a non-negligible role.

In the next section, we will start by establishing a connection between the phenotype-only and the gradient models (Section 5.2). This will enable us to show that the phenotypic clusters observed in Figures 2–4 arise for the same reasons. Next, we will aim at understanding those reasons: we will demonstrate that the emergence of clusters is not due to Turing patterns (Section 5.3), but to demographic noise, and we will present some quantitative results on admissible spacings (Section 5.4). Lastly, in Section 5.5, we will consider different rates of recombination and sex, and will examine their influence on the emergence of phenotypic clusters.

### 5.2 Connection between the two models

Let us show how the gradient and the phenotype-only models are connected. We do this by separating the ‘ecological’(fast) and the ‘evolutionary’(slow) timescales in the gradient model. To this aim, we work under the following conditions: (*i*) the persistence criterion (4) holds, i.e. the population can survive; (*ii*) the effect of mutations *m* is small, and can be replaced with 0 in the fast timescale; (*iii*) the solution of the gradient equation (3) converges smoothly to an equilibrium; and (*iv*) *ρ*(*t, x*) does not vary much in comparison with (*z* −*bx*)^2^/2*V*_*s*_, so that we can write

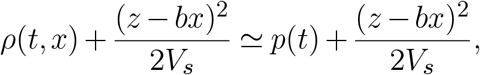

for some function *p*(*t*) independent of *x*.

Assumptions (*i*) and (*ii*) only depend on the parameters of the model and are easy to verify. Assumption (*iii*) is natural and is supported by simulations, although it cannot be taken *a priori* for granted (see in particular Nadin *et al*. 2013, where oscillations in time occur in a similar model with nonlocal time-dependence). Simulations suggest that the approximation made in Assumption (*iv*) is sufficiently good that our results hold.

We let *u*(*t, x, z*) be the solution of (3). We write, for *t* ⩾ 0, and, 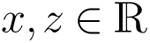,

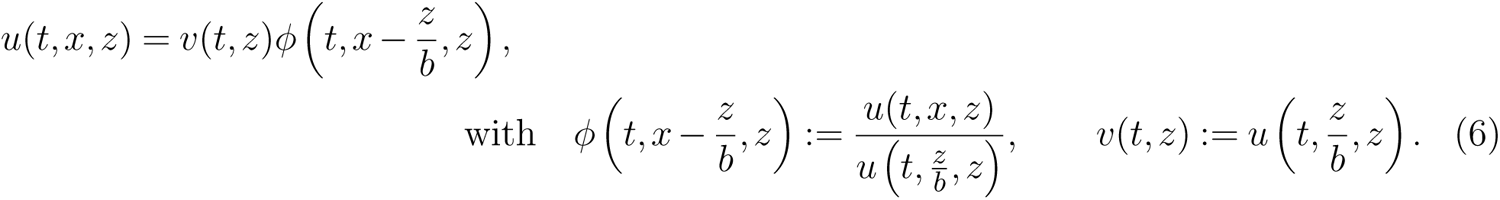

In words, *v*(*t, z*) is the density of the population carrying phenotype *z* at the optimal geographical position associated with *z*, and *x* ↦ *ϕ*(*t, x*− *z*/*b, z*) is the spatial distribution of the population around the optimal geographical position associated with *z*.

The function *ϕ* contains the dynamics of *u* on the (fast) ecological timescale, while the function *v* contains the dynamics of *u* on the (slow) evolutionary timescale. We will show that under conditions (*i*)–(*iv*), letting 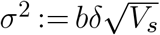, the function *ϕ* is close to a Gaussian equilibrium (independently of *z*): 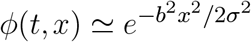. Using this, we will show the main result of this section:

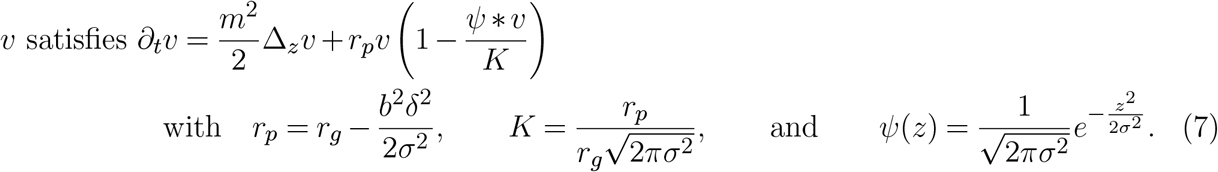

Observe that *r*_*p*_ is positive due to the persistence criterion (4), and that the equation satisfied by *v* is almost identical to (1), except for the carrying capacity *K* (which has no effect on the qualitative behavior of the solution).

First, using (6), and recalling that *u* satisfies (3) with *m* ≃ 0 (Assumption (*ii*)), we have:

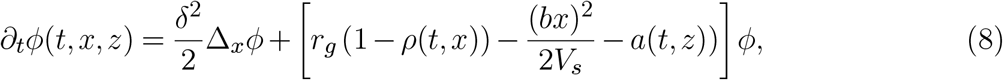

with *a*(*t, z*) := *∂*_*t*_*v*(*t, z*)/*v*(*t, z*). Details are provided in Appendix B. Next, owing to Assumptions (*iii*) (*ρ*(*t, x*) = *p*(*t*)) and (*iv*) (smooth convergence to an equilibrium), we must have, as *t* → +∞,

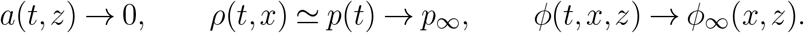

In Appendix B, we show that necessarily 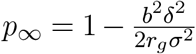 and 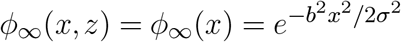, with *σ*^2^ defined above. Note that *ϕ*_∞_ does not depend on *t* and *z*. Last, replacing *ϕ*(*t, x, z*) with *ϕ*_∞_ in (6), and after a few computations detailed in Appendix B, we obtain that (7) holds.

Hence, we proved that the phenotype-only and the gradient models could be related to each other via the approximate relation

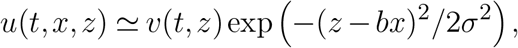

where *u* solves the gradient equation, *v* solves the phenotype-only equation (where the competition kernel *ψ* is a Gaussian with variance *σ*^2^), and 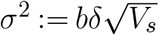. Therefore, phenotypic clusters appear in *u* if, and only if phenotypic clusters appear in *v*. Keeping this in mind in the sequel, we will study the phenotype-only model with a Gaussian competition kernel, and then extend those results to the gradient model.

### 5.3 Turing instability cannot explain the emergence of evenly-spaced patterns

The behaviour of the phenotype-only equation (1) is complex and has raised a wide interest in the literature (see e.g., Genieys *et al*. 2006; Berestycki *et al*. 2009; Hamel & Ryzhik 2014). The reason of such an interest and of such a complexity is that for some sets of parameters, Turing patterns arise in the solutions of (1). In this section, we start by giving an idea of why such patterns could arise. Next, we will show that this phenomenon is *not* the cause of the patterns that we observe in our simulations.

#### Turing instability

To start with, note that *V*_*eq*_ ≡ 1 is an equilibrium state of (1). Let 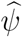 be the Fourier transform of *ψ*, defined by

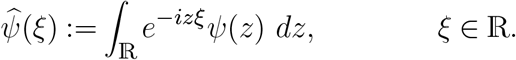

In Genieys *et al*. (2006), the authors showed by heuristic arguments and simulations that if there exists *ξ* ∈ ℝ such that 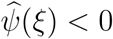, then, for sufficiently small mutation variance *m* and when started from a perturbation

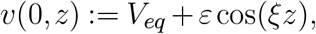

the perturbation is amplified for some small *ε* > 0. In such cases, evenly-spaced patterns with spacing 2*π*/*ξ* are expected to appear in the solution. This means that the equilibrium *V*_*eq*_ ≡ 1 is unstable, and the evenly-spaced patterns are called *Turing patterns*. For example, simulations in Genieys *et al*. (2006) show that when *ψ*(*z*) = *ψ*_*A*_(*z*) with 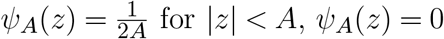 for |*z*| *< A, ψ*_*A*_(*z*) = 0 for |*z*| ⩾ *A*, then patterns appear for large values of *A*, for which 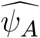 takes negative values. Rigorous works on Equation (1) have followed, and proved the role of the sign of the Fourier transform of the competition kernel *ψ* (Berestycki *et al*. 2009; Fang & Zhao 2011; Hamel & Ryzhik 2014). Conversely, when 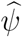 does not change sign, Turing patterns do not appear: the uniform equilibrium is stable and all perturbations of the kind mentioned above are damped.

#### In our simulations, there are no Turing instabilities

In our case, the competition kernel *ψ* is a Gaussian with variance *σ*^2^ (by design for the phenotype-only model, and by the above analysis for the gradient model), and therefore has a Fourier transform 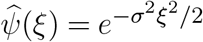. In particular, 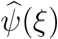 is positive for all *ξ*. As a consequence, the equilibrium *V*_*eq*_ ≡ 1 is stable, and we are *not* in the situation where evenly-spaced patterns arise (in particular, there are no non-constant steady states to (1), see Berestycki *et al*. 2009, Theorem 1.2).

Figure 2 confirms this: in the phenotype-only model, no patterns appear for large populations, which implies that patterns cannot be observed through the deterministic point of view of Turing instability. Figure 4 shows that the same conclusion can be reached for the gradient model.

#### About the patterns observed in Doebeli & Dieckmann (2003)

Patterns similar to those of the left panel in Figure 4, were observed by Doebeli & Dieckmann (2003) in a model very close to the gradient model (Equation (3)). The only difference is in the competition term *ρ*(*t, x*), which is replaced with

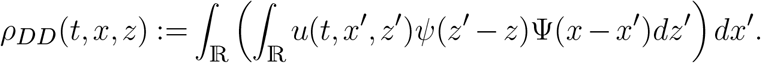

Namely, as in the phenotype-only model, the competition between phenotypes *z* and *z*′ depends on the distance between those two phenotypes via the kernel *ψ*. Contrarily to our model, individuals present in different positions interact via the *geographical* competition kernel Ψ. With this transformation, the authors observed the formation of phenotypic clusters.

Polechová & Barton (2005) argued that these patterns are merely a consequence of boundary effects. However, this interpretation appears insufficient, since the same patterns are also observed in simulations carried out on domains explicitly devoid of boundaries. A different explanation was put forward by Leimar *et al*. (2008), where it was shown that these patterns may arise as a consequence of Turing instability for a certain class of geographical competition kernels Ψ (i.e., the competition kernels whose Fourier transform changes sign). Yet the geographical competition kernel considered in Doebeli & Dieckmann (2003), is a Gaussian and therefore does not belong to this class. In other words, the results of Leimar *et al*. (2008) do not directly apply to the setting under consideration in Doebeli & Dieckmann (2003).

In the next section, we provide an alternative explanation that is independent of both of the above interpretations. We show, indeed, that the patterns observed in Doebeli & Dieck- mann (2003) emerge naturally as a consequence of demographic noise. We attached a Jupyter Notebook to the supplementary materials with simulations specific to the Doebeli-Dieckmann model, that support the role of demographic noise.

### 5.4 Demographic noise explains the emergence of evenly-spaced patterns

We claim that the emergence of patterns that are observed in our simulations is due to instability in tightly packed species at low population levels. These patterns are therefore a purely stochastic phenomenon. In order to gain insight into the effects of stochasticity, and to quantify the tightest possible packing, we consider an approximation of (2) which results in an equation similar to that of May & MacArthur (1972). This approximation provides a sufficiently simple framework to study stable spacings, and fits our simulations very well.

#### May-MacArthur approximation

Consider the phenotype-only model (Equation (2)) with mutation variance *m*^2^ = 0 and an initial condition of the form

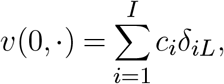

where, for *i* ∈ {1,…, *I*}, *δ*_*iL*_ is a Dirac mass at *iL* and *c*_*i*_ > 0 is a given coefficient. Here, *I* is the total number of coexisting clusters and is assumed to be large. We work in the periodic setting, so that the Dirac mass at position *I* is at a distance *L* from the Dirac mass at position *I* − 1 and from the Dirac mass at position 1. Here, *L* > 0 is the distance between two adjacent clusters *i* and *i* + 1 at the initial time. The question is: *if there is a distance L between two adjacent clusters, is the population stable?*

With *m* = 0 and such an initial condition, (2) can be understood as a (large) system of SDEs. For *i, j* ∈ {1,…, *I*}, let *d*(*i, j*) := min(|*i*− *j*|, *I* − |*i*− *j*|) be the distance between *i* and *j* on the torus, and let *a*_*i,j*_ := *ψ*(*L*× *d*(*i, j*)) be the strength of the interaction between species *i* and *j*. Then, for all *t* ⩾ 0, *v*(*t*, ·) can be written in the form

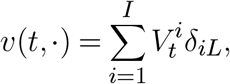

where, for each *i* ∈ {1,…, *I*}, 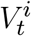 solves the equation:

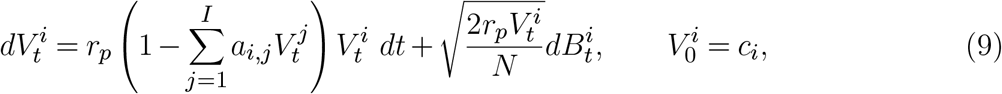

for some independent Brownian motions 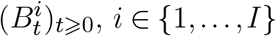.

For *N* = +∞, there is no noise and we obtain exactly the equation studied by May & MacArthur (1972). In this case, Equation (9) is deterministic and has an equilibrium that satisfies

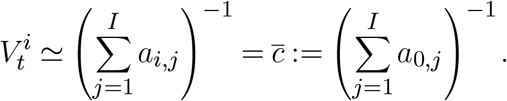

We shall be interested in the stability, on a timescale of order *O*(*N*), of the equilibrium state 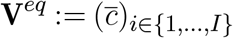.

#### For which values of *L* is the equilibrium of (9) stable?

From now on, we fix *σ* > 0 and we define *η* := *L*/*σ*, where *L* is the space between two adjacent clusters. Let 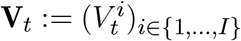 denote the population vector of the solution of (9). Assume that **V** is a small perturbation of the equilibrium, that is, write 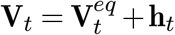 for small **h**_*t*_. We show in Appendix C that as long as **h** is small, it solves the equation

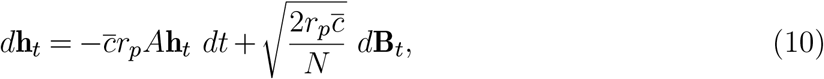

where *A* is the (*I* × *I*)-matrix with coefficients (*a*_*i,j*_)_*i,j*∈{1,…,*I*}_, and **B**_*t*_ is a Brownian vector of size *I*.

Accelerating time by a factor 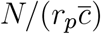 gives:

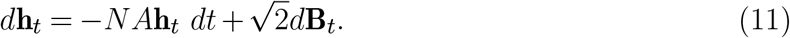

In our simulations, we neglect the dependence of 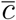 in *η* (on the range of values of *η* that appear, this is inconsequential). Letting *λ*[*η*] be the principal eigenvalue of the matrix *A*, the dynamics of Equation (11) looks like

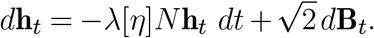

In this new equation, the process **h**_*t*_ is submitted to two opposite phenomena: the term 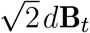 makes **h**_*t*_ evolve randomly on a timescale of order 1, while −*λ*[*η*]*N* **h**_*t*_ *dt* brings **h**_*t*_ closer to 0 on a timescale of order 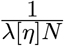. We conclude that a sufficient condition for the stability of (9) up to a time horizon of the order *N* should be:

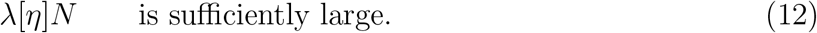

As *I* → +∞, the limiting value of the principal eigenvalue *λ*[*η*] is given explicitly in May & MacArthur (1972). With our notations, the limiting value is:

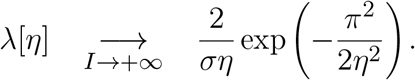

We take *I* large so that equality can reasonably be assumed to hold.

#### Connection between *N* and the observed distances between adjacent clusters

Let us study the condition (12) in order to understand the connection between *N* and the observed spacing. Let us start by rewriting (12) as

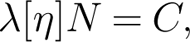

for some fixed constant *C* > 0 which does not depend on *N* . After some computations (see Appendix C), we find that for *η* ∈ (0, *π*) and 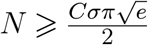, the above condition becomes:

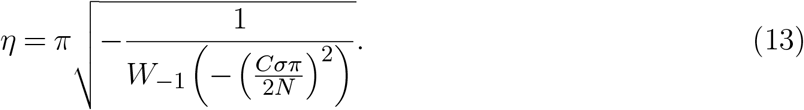

Here, *W*_−1_ denotes the branch of index −1 of the Lambert *W* function. More precisely, for *a* ∈ [−1/*e*, 0), *x* = *W*_−1_(*a*) is the smallest real solution of *xe*^−*x*^ = *a*.

Despite the rough approximations that we have performed, simulations show a good agreement between the observed spacing and the above prediction: namely, Figure 5, left, shows that the minimal stable spacing is very well approximated by (13). In order to be as close as possible to our heuristics, the simulations of Figure 5, left (and only those), were run with the Gaussian approximation and with *m* = 0, i.e., without mutations.

**Figure 5.**
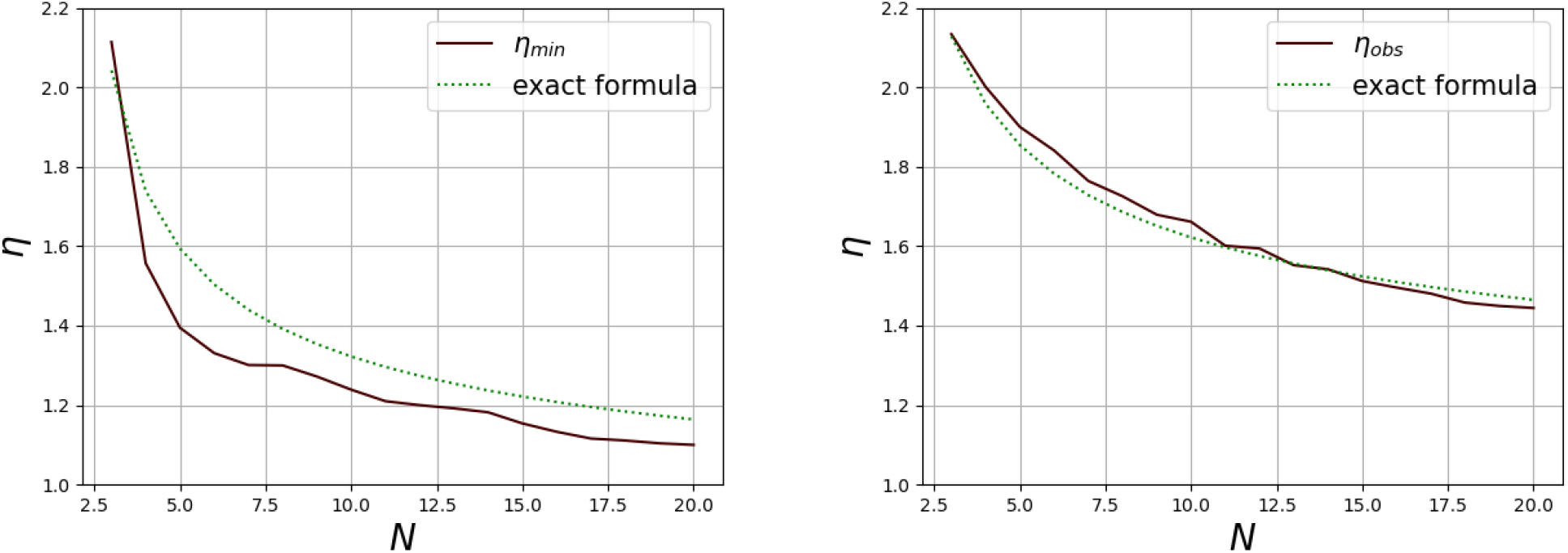
Phenotype-only model: Simulated values of *η*_obs_ := *L*/*σ* are displayed against *N*, and compared with the relation (14) (“exact formula”). **Left: Minimal stable spacing (Gaussian and no-mutation approximation**, *m* = 0). Each value of observed *η*_min_ is computed by taking *L*_min_/*σ*, where *L*_min_ is the average of the observed minimal stable spacing over 100 replicates. Equation (14) is displayed with *C* = 0.09 and *γ* = 1. **Right: Spacing between spontaneous clusters (Original model**, *m* = 0.01). Each value of observed *η*_obs_ is computed by taking *L*_obs_/*σ*, where *L*_obs_ is the average of the observed spacing over 200 replicates, when started from a uniform population. Equation (14) is displayed with *C* = 0.05 and *γ* = 1.37. **Both figures**. The competition kernel *ψ* is a Gaussian with variance *σ*^2^ = 100. The time at which *η* was calculated is *t* = *t*_0_*N*, with *t*_0_ = 1000. The parameter *N* takes 18 values ranging from 3 to 20.

We next focused on the original model without the Gaussian approximation and with a positive mutation rate. We estimated the observed spacing when started from a uniformly distributed population, i.e., the spacing between clusters that spontaneously arise. To this aim, we tried to fit our observations with a variant of Equation (13). Namely, we observed that (13) holds up to a multiplication by some *γ*, namely:

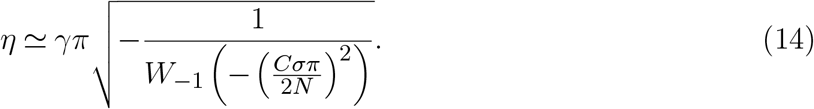

See Figures 5, right. Observe that at equilibrium, clusters can be more tightly packed if the initial population is not uniformly distributed (as in Figure 5, right), but rather already made up of sufficiently tightly packed clusters (as in Figure 5, left). Below, we will focus in more details on the effect of the initial spacing on the observed spacing at equilibrium.

For small 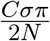, we can use the asymptotic expansion of *W*_−1_ near 0 and find that

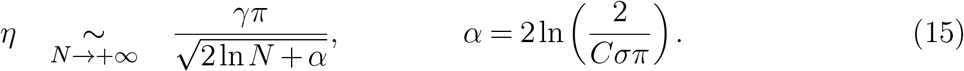

See Appendix C. This shows that the observed spacing does decrease with *N*, but extremely slowly.

#### The effect of *σ*

From (15), one finds that the constant *α* depends only logarithmically on *σ*, and therefore, that *η* is almost independent of *σ*, that is, that *L* ≃ *ησ*. Figure 6 suggests that at least for small values of *σ*, the above expression holds, up to some error term, i.e., one has in fact *L* = *ησ* + *b* for some *b* ∈ ℝ (also depending on *N*).

**Figure 6.**
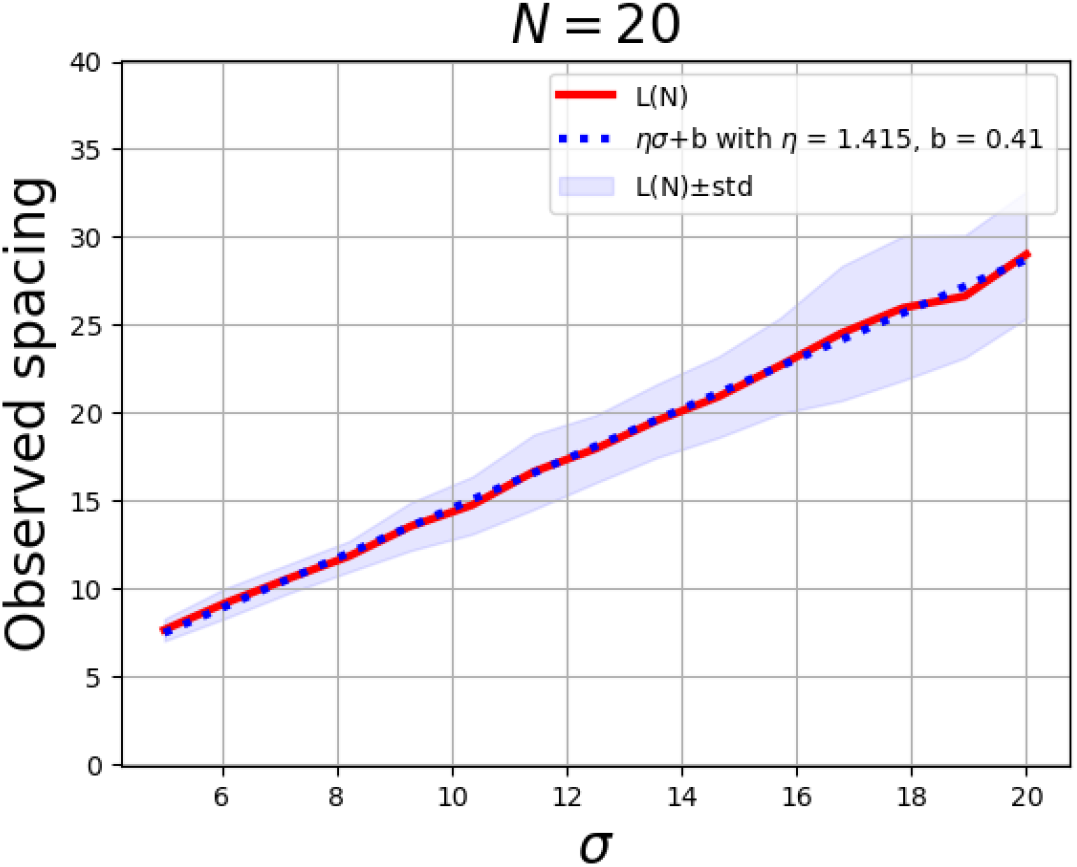
Observed spacing *L* with respect to the variance *σ*^2^. The affine best fit is plotted in dotted blue. The time at which *η* was calculated is *t* = *t*_0_*N*, with *t*_0_ = 1000. Here, *N* = 20, *σ* takes 15 values between 5 and 20, and 50 replicates were run for each value of *σ*. The other parameters are the same as for Figure 5, right.

#### Consequence on the gradient model

Let us now extend these observations to the gradient model. Via the correspondence with the phenotype-only model, we tried to fit the observed spacing between spontaneously emerging clusters in the gradient model with Equation (14). Figure 7 shows that the agreement is, again, very good.

**Figure 7.**
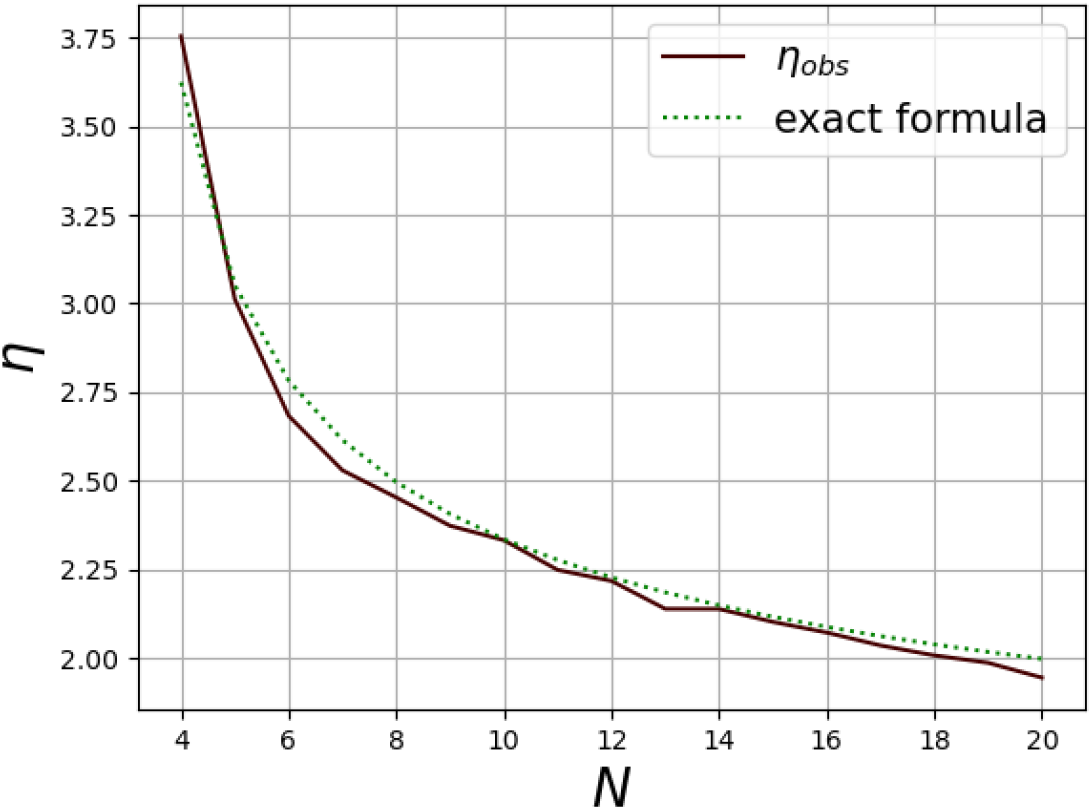
Gradient model: Simulated values of *η*_obs_ := *L*/*σ* are displayed against *N*, and compared with the relation (14) (“exact formula”). Each value of observed *η*_obs_ is computed by taking *L*_obs_/*σ*, where *L*_obs_ is the average of the observed spacing over 200 replicates. The time at which *η* was calculated is *t* = 50*N* . The parameter *N* takes 17 values ranging from 4 to 20. The other parameters are as in Figure 4. On this figure, we have *γ* = 1.6 and *C* = 0.27.

#### Dependency on the initial spacing

Next, we start the simulations for the phenotype-only model from already clearly defined phenotypic clusters, with different spacings *L*_0_ ranging from 1 to 20. We identified two discontinuous transitions in the connection between population size, the initial distances between clusters, and the distances between clusters at equilibrium (Figure 8).

**Figure 8.**
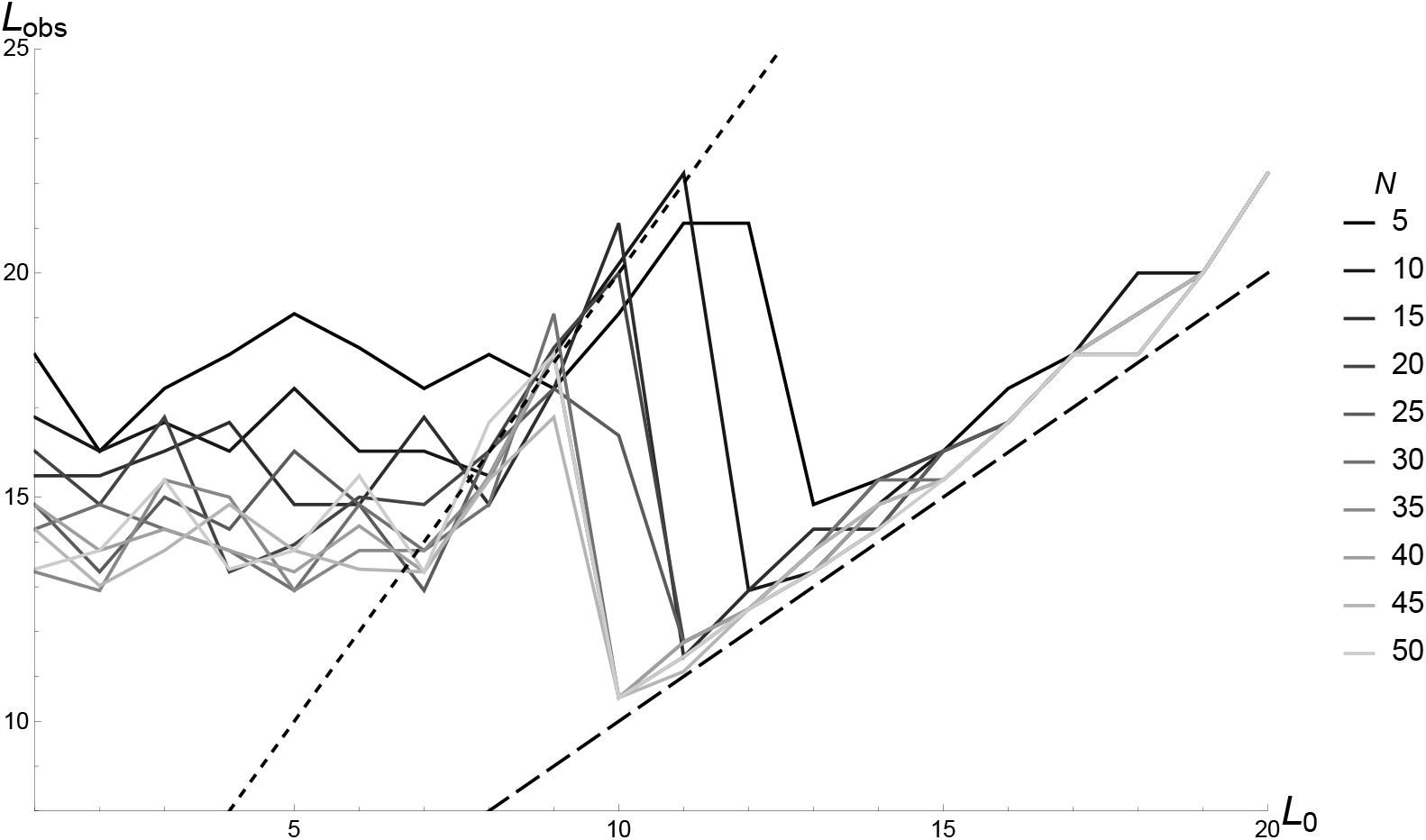
Illustration of the three regimes in the connection between population size, the initial spacings between clusters, and the spacings between clusters at equilibrium. The horizontal axis corresponds to *L*_0_ (spacing at the beginning of the simulation) and the vertical axis to *L*_obs_ (average of the observed spacing over 50 replicates, with parameter *σ* = 10). The color gradient correspond to ten different values of the carrying capacity *N* used in the phenotype-only model. The two dashed diagonals correspond to *y* = 2*x* (left) and *y* = *x* (right), fitting well with the equilibrium spacings in the regimes *L*_−_(*N*) ⩽ *L*_0_ ⩽ *L*_+_(*N*) and *L*_0_ ⩾ *L*_+_(*N*), respectively.

More precisely, there are three regimes: (*i*) small values of the initial spacing *L*_0_ (*L*_0_ ⩽ *L*_−_(*N*)), for which the spacing at equilibrium, *L*_obs_, is the same as for a uniform initial population; (*ii*) intermediate values of the initial spacing *L*_0_ (*L*_−_(*N*) ⩽ *L*_0_ ⩽ *L*_+_(*N*)), for which *L*_obs_ = 2*L*_0_; and (*iii*) large values of the initial spacing *L*_0_ (*L*_0_ ⩾ *L*_+_(*N*)), for which *L*_obs_ = *L*_0_. The transitions *L*_−_(*N*) and *L*_+_(*N*) between the regimes decrease when *N* increases.

In the regime *L*_0_ ⩽ *L*_−_(*N*), the initial spacing is so small that it is not felt by the population dynamics, i.e., the long-term observations are the same as when starting from an initial population without phenotypic clusters. In the regime *L*_0_ ⩾ *L*_+_(*N*), the initial spacing is already stable, and therefore persists in time. In the intermediate regime *L*_−_(*N*) ⩽ *L*_0_ ⩽ *L*_+_(*N*), the initial spacing is unstable, but not sufficiently small to be equivalent to an initial uniform population. In these simulations, we observe that roughly one cluster out of two disappears, leading to *L*_obs_ = 2*L*_0_.

In particular, the above remarks imply that the relationship between the spacing at the initial time and the spacing at equilibrium is not monotonic. We also note that *L*_+_(*N*) is strictly smaller than the spacing at equilibrium when starting from a population without phenotypic clusters. As a consequence, starting from a uniform population without phenotypic clusters does not necessarily result in the tightest packing; in fact, the tightest packing is obtained with initial spacing values slightly larger than *L*_+_(*N*).

### 5.5 Simulations of the model with sexual reproduction

In this section, we are interested in exploring what happens in the gradient model when we relax the assumption of total clonality. Namely, we progressively increase the effective rate of recombination (*r*_eff_) in the parameter values where we expect phenotypic clustering to occur (as in Figure 4). To investigate the effect of *r*_eff_, we varied the rate of sex (*π*_sex_), as well as the average number of crossovers (*R*) per meiotic event. Those two parameters affects the effective recombination rate *r*.

As seen in Figure 9, the stair steps are less visible in this new setting, making the use of the spacing *L* between consecutive phenotypic clusters less appropriate. To study the effect of *r*_eff_ on phenotypic clusters, we turned to the amplitude and the period of the main oscillations observed in the phenotypic variance (*V*_*z*_) and the deviation of the mean phenotype from the local phenotypic optimum 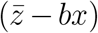. By doing so, we observe that the effective recombination rate has a very strong effect on the amplitude, but no effect on the period of the main oscillations (Figures 10 for *V*_*z*_ and S1 for 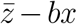).

**Figure 9.**
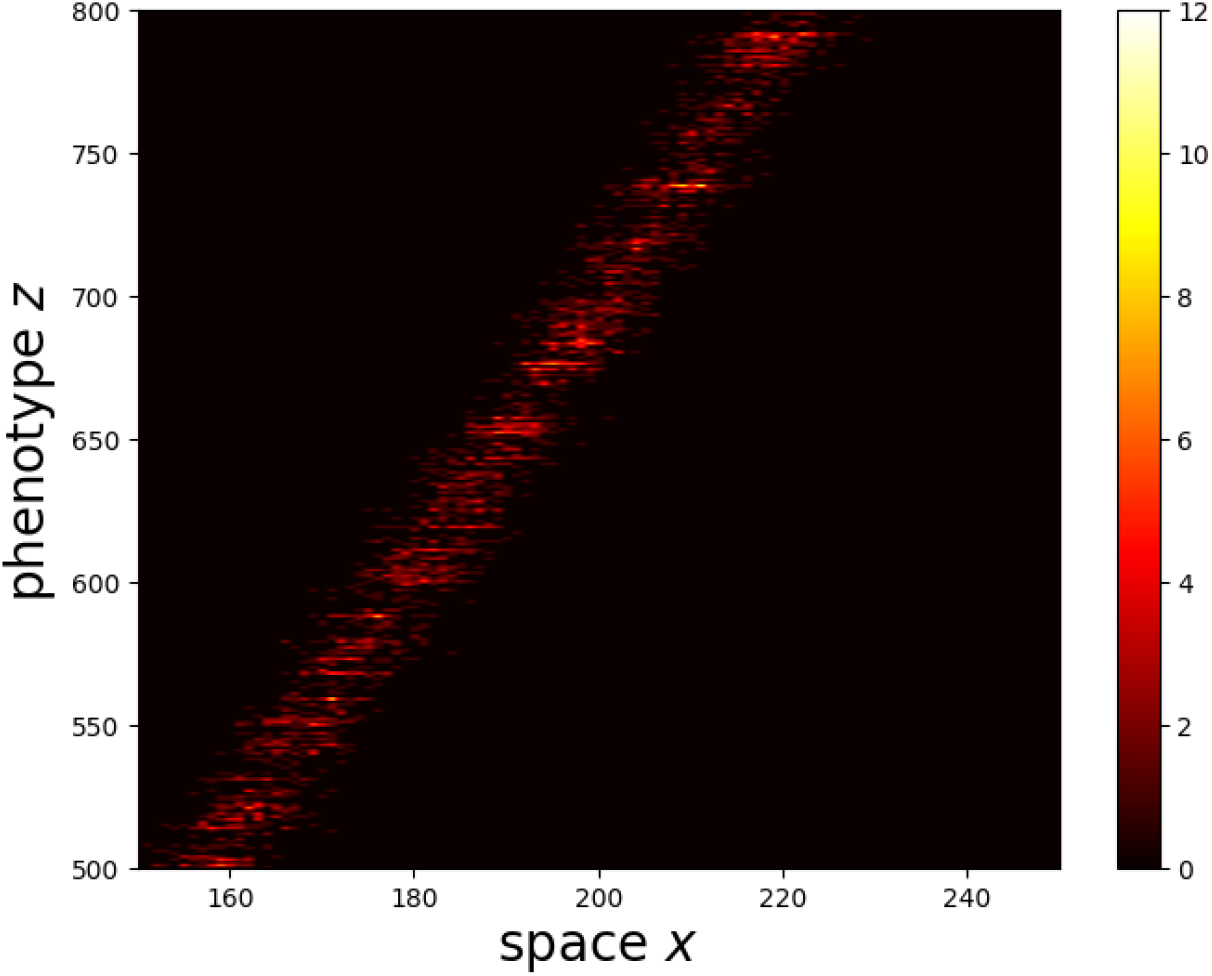
Output of the gradient model with sexual reproduction at the end of the simulation (time *t* = 50, 000). The horizontal axis corresponds to the geographical space *x* (here between demes 150 and 250), and the vertical axis corresponds to the phenotypic space *z*. The color gradient corresponds to the density of individuals having phenotype *z* in position *x*. The slope of the environmental gradient is set to *b* = 1.831, thus the scales differ between the vertical and horizontal axes. The values of the parameters impacting the effective recombination rate are: *R* = 100 and *π*_sex_ = 0.1, leading to *r*_eff_ = 0.05.

**Figure 10.**
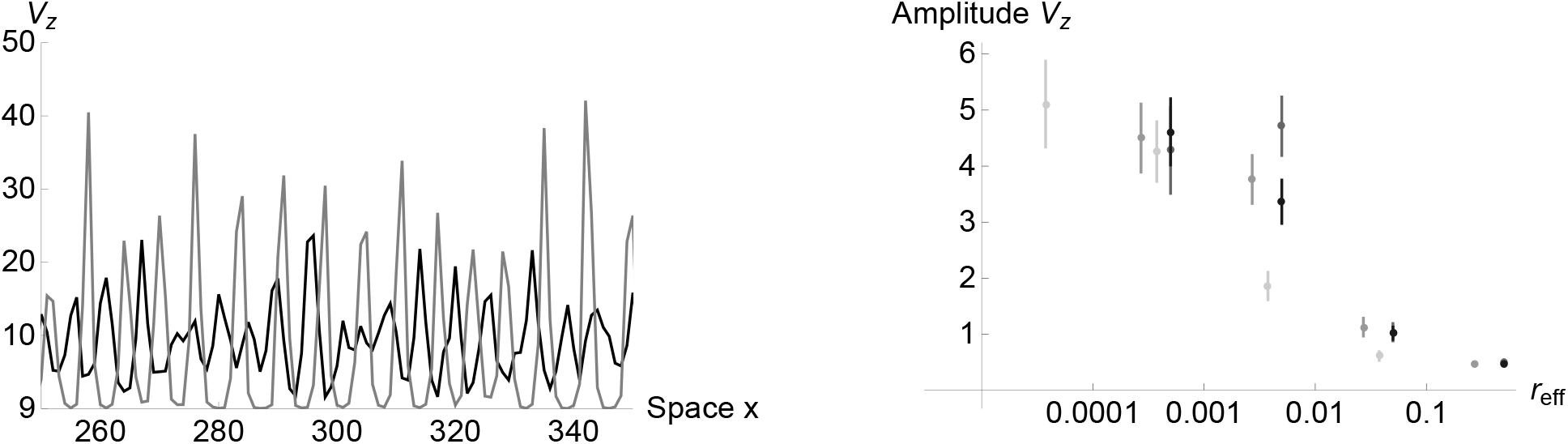
**Left:** Phenotypic variance *V*_*z*_ measured in each deme *x*, here shown for *x* ∈ [250, 350]. The gray curve corresponds to the results obtained for *r*_eff_ = 4.10^−6^ (*R* = 0.1, and *π*_sex_ = 10^−3^); and the black curve to the one obtained for *r*_eff_ = 4.10^−3^ (*R* = 1 and *π*_sex_ = 10^−1^). **Right:** Amplitude of the fluctuations observed in the phenotypic variance *V*_*z*_ after selection, as a function of the effective recombination rate *r*_eff_ (here on a log scale). The effective recombination rate *r*_eff_ is defined as the product of the recombination rate and the rate of sex. The different shades correspond to different sets of *R* and *π*_sex_. The values of the remaining parameters are the same across simulations and correspond to: *N* = 100, *s* = 0.005, *V*_*s*_ = 12.488, *α* = 0.255, *b* = 1.831, *σ* = 2.591 and *L* = 2561. The mean and standard error were measured across ten replicates.

Indeed, as can be seen on the left of Figure 10, a 1000-fold difference in *r*_eff_ mainly leads to a difference in the amplitude of the fluctuations (Figures 10 and S1) while keeping the period of the main oscillation almost unchanged. This leads to a weak signal of the effect of *r*_eff_ on the observed period (Figure S2).

## 6 Discussion

Understanding how and why a finite number of phenotypically distinct species or populations emerge along a seemingly continuous resource gradient has been a central question in theoretical ecology, and is closely related to the principle of competitive exclusion (May & MacArthur 1972). In mathematical models, competitive exclusion is commonly represented through nonlocal competition in the phenotype space, an approach initiated in reaction-diffusion models by Britton (1989). These models can produce separated groups in the phenotype space, often referred to as *phenotypic clusters*, which may correspond to distinct coexisting ecological types or incipient species.

Most previous studies on phenotypic clustering have focused on deterministic mechanisms. In particular, clustering has often been interpreted as the result of Turing instability arising from the shape of the competition kernel (Sasaki 1997; Genieys *et al*. 2006; Leimar *et al*. 2008). In contrast, our results show that phenotypic clustering can emerge from demographic stochasticity alone, even in regimes where deterministic Turing instability is absent.

We first highlighted this mechanism in a one-dimensional model that we dubbed”phenotypeonly”. In this framework, we incorporated the effect of mutation and competition between individuals. By considering a Gaussian competition kernel, we did not place ourselves in the context where Turing instability is expected. Demographic noise was incorporated by accounting for random fluctuations associated with finite population sizes. Our results have shown that a sufficient amount of demographic noise could lead to evenly-spaced phenotypic clusters. These clusters were indeed only observed for small population sizes, corresponding to stronger demographic fluctuations (Figure 2). Moreover, these clusters distinctly emerged very rapidly in time (Figure 3). Similar stochastic clustering was previously reported by Rogers *et al*. (2012), with a focus on different scalings of demographic noise. Here, we extended the results in the strong noise regime by connecting stochastic clustering to environmental gradients, and by deriving an explicit relationship between spatial structure, effective competition in phenotype space, and demographic noise.

We subsequently considered a model, referred to as the “gradient model”, involving a linear geographic space where each position is associated with an optimal phenotype upon which the phenotypic trait is submitted to stabilizing selection and to a density-dependent component. This is a common framework in population and quantitative genetics (Felsenstein 1977; Kirkpatrick & Barton 1997; Polechová & Barton 2015; Fouqueau & Roze 2021). As in the phenotype-only model, we observed the formation of phenotypic clusters in the gradient model (Figure 4). A central result of our work is that the gradient model can be connected to the phenotype-only model by separating the fast and slow timescales (Section 5.2). More precisely, while individuals rapidly redistribute around the geographical location where their phenotype is locally optimal, the total density associated with each phenotype evolves more slowly through competition and demographic fluctuations. Using this connection, we were able to show that under certain conditions, the geographical distribution of individuals carrying a given phenotype converges to a Gaussian centered around their optimal geographical position. Importantly, the correspondence between the phenotype-only and the gradient models enabled us to show that the variance in the geographic distribution can be expressed in terms of the slope of the environmental gradient (*b*), dispersal (*δ*) and stabilizing selection 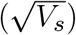.

It is important to stress that the emergence of phenotypic clusters in the gradient model cannot be attributed to deterministic Turing instabilities. The fact that the effective competition kernel is Gaussian, and thus Turing-stable, is a consequence of the quadratic shape of the selection term, 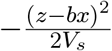. However, it is unclear whether this outcome is robust to changes in the selection term. In fact, we have been unable to find a general selection term that leads to a Turing-unstable competition kernel *ψ* (the selection term is of the form *R*(|*z* −*bx*|), for some general decreasing function *R*, and *ψ* is the only positive solution of the eigenvalue problem (16)). This suggests that for the gradient model with only within-deme competition, Turing stability is not a special limiting case, contrarily to the phenotype-only model (Barabás & Meszéna 2009; Pigolotti *et al*. 2010; Leimar *et al*. 2013).

We have mentioned until now that the emergence of phenotypic clusters depends on stochasticity. Once these phenotypic clusters have appeared, it is only natural to wonder how far apart they are. We have explored this question by adapting the results of May & MacArthur (1972) to our framework. We have shown how the distances between successive clusters at equilibrium depend on population size, in both the phenotype-only model (Figure 5) and the gradient model (Figures 6 and 7). We emphasize that the relations displayed in Figures 5–7 are only valid for ranges of parameters for which the environment is effectively “almost infinite”. Out of these ranges of parameters, the formation of phenotypic clusters is driven more by the boundedness of the simulated environment than by the instability of too tight packing, invalidating the heuristics of Section 5.4. In addition to the effect of population size, we have found that the initial spacing between phenotypic clusters additionally influences the equilibrium spacing, with some discontinuous and non-monotonic transitions (Figure 8). This outcome is due, on the one hand, to the high level of stochasticity, which prevents dense packing. On the other hand, it is also due to the small effect of mutations (on the value of the phenotype), which prevents denser packing structures from emerging.

Finally, we relaxed the assumption of complete clonality by introducing sexual reproduction and recombination, whose combined effects determine the effective recombination rate (Roze 2014). This extension is important because phenotypic clustering in clonal models is sometimes interpreted as a form of sympatric speciation (e.g., Dieckmann & Doebeli 1999; Rogers *et al*. 2012). However, in the absence of sexual reproduction, the biological meaning of the term “species” remains ambiguous, since reproductive isolation cannot emerge (Kulmuni *et al*. 2020). We have shown that phenotypic clusters quickly vanish as the effective recombination rate increases (Figures 10 and S1). In this context, phenotypic clusters should rather be interpreted as coexisting ecological or phenotypic types than as fully distinct species.

## Acknowledgments

This work stems from the *Probability meets Biology* workshop that was held in November 2024 in Berlin. We wish to thank the organizers of this workshop. N.B. received funding from the FWF Project PAT3816823 “Waves in Population Genetics”. L.F. acknowledges the support of the NOMIS-ISTA Fellowship.

## Data availability

We performed a number of simulations of our models. The models with asexual reproduction where simulated in Python; the corresponding Jupyter notebooks are available at doi: 10.5281/zenodo.21505286. The model with sexual reproduction was simulated in C++; the code is available at doi: 10.5281/zenodo.4643637 (Fouqueau & Roze 2021).

## A Supplementary Figures

**Figure S1.**
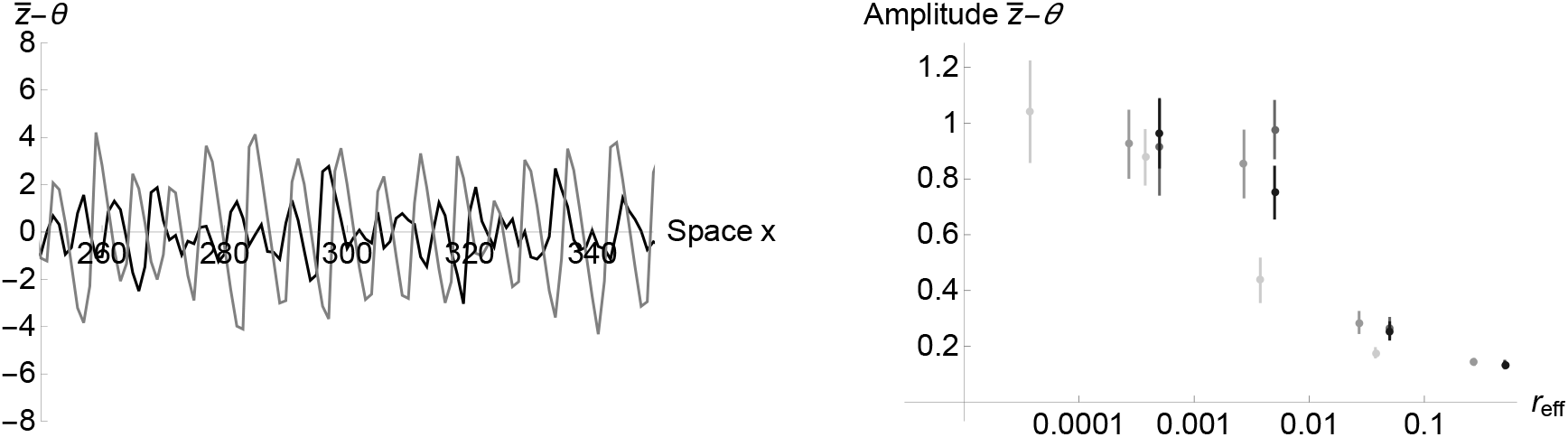
**Left:** Deviation of the mean phenotype from the local phenotypic optima 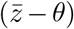, measured in each deme *x*, here shown for *x* ∈ [250, 350]. The gray curve corresponds to the results obtained for *r*_eff_ = 4.10^−6^, *R* = 0.1, and *π*_sex_ = 10^−3^; and the black curve to *r*_eff_ = 4.10^−3^ obtained for *R* = 1 and *π*_sex_ = 10^−1^. **Right:** Amplitude of the fluctuations observed in 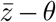 after selection, as a function of the effective recombination rate *r*_eff_ (here on a log scale). The different shades correspond to different sets of values of *R* and *π*_sex_ leading to similar values of *r*_eff_. The values of the remaining parameters are the same across simulations and for both figures, and correspond to: *N* = 100, *s* = 0.005, *V*_*s*_ = 12.488, *α* = 0.255, *b* = 1.831, *σ* = 2.591 and *L* = 2561. The mean and standard error were measured across ten replicates.

**Figure S2.**
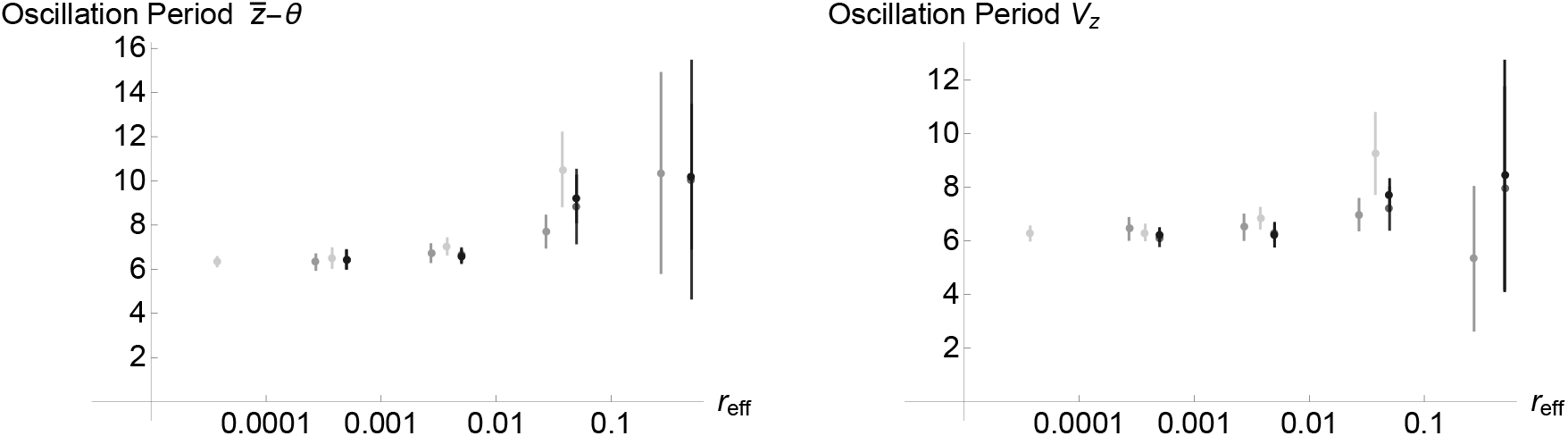
Period of the main oscillation of the Fourier transform for the deviation of the mean phenotype from the local phenotypic optimum (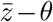, **left**) and of the phenotypic variance (*V*_*z*_, **right**) as a function of the effective recombination rate *r*_eff_. The values of the remaining parameters are the same as in Figure S1

## B Connection between the PO and the gradient models

### Ecological timescale: Proof that *ϕ* **given by** (6) **satisfies** (8)

We first work in the (fast) ecological timescale, where we assume that the effect of mutations can be neglected (Assumption (*ii*)), i.e. we take *m* = 0. Let *ϕ* be defined by 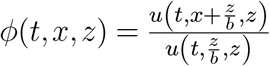, where *u* is a solution of (3) with *m* = 0. Let 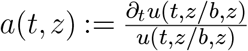. We have:

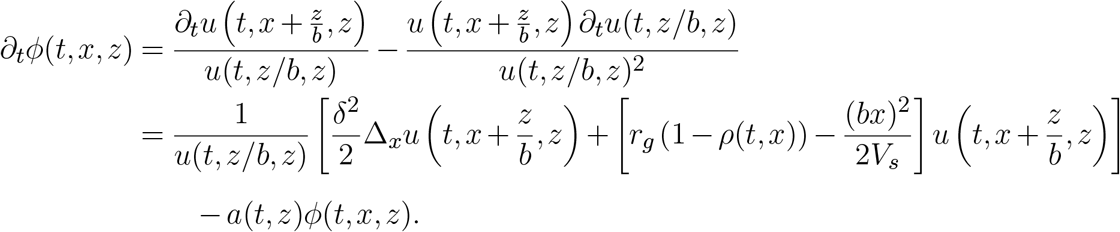

Now, we note that 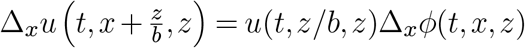, and we obtain:

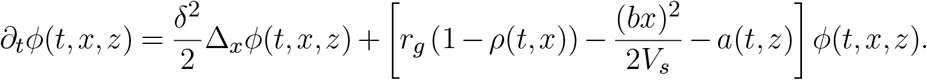

This is precisely (8).

### Analysis of *ϕ*_∞_

Owing to Assumption (*iii*) (*ρ*(*t, x*) = *p*(*t*)) and Assumption (*iv*) (smooth convergence to an equilibrium), we must have, as *t* → +∞,

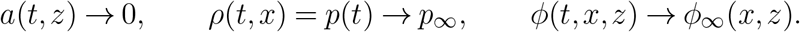

Let *L* be the operator defined by 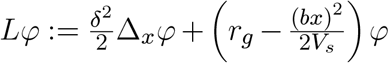. Taking *t* → +∞ in (8), one obtains that that *ϕ*_∞_ is a positive solution of

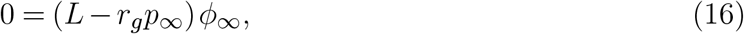

so *ϕ*_∞_ is a positive eigenfunction of *L* associated with the eigenvalue *r*_*g*_*p*_∞_. Recall also that *ϕ*(*t*, 0, *z*) = 1. Let us now show that it allows us to characterize *ϕ*_∞_ and *p*_∞_ at the same time.

Consider, for some *σ*^2^ > 0, the Gaussian function 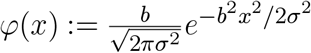. Computations show that

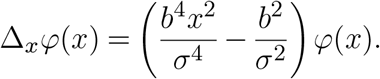

Therefore,

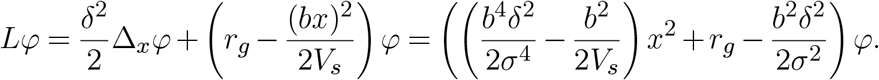

Now, taking 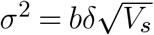, we have

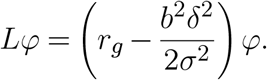

Hence, *φ* is a positive eigenfunction of *L*, associated with the eigenvalue 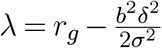. The operator *L* : *H*^2^(ℝ) ⊂ *L*^2^(ℝ) → *L*^2^(ℝ) has compact resolvents (owing to the fact that the potential term 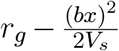 is confining). Hence, by the Krein-Rutman theory, any positive eigenfunction of *L* is a (positive) multiple of *φ*, and is associated with the eigenvalue *λ*. We conclude that *ϕ*_∞_ is a positive multiple of *φ* and that *r*_*g*_*p*_∞_ = *λ*. Pointing out that *ϕ*(*t*, 0, *z*) = 1 for each *t* > 0 and *z* ∈ ℝ, we obtain that *ϕ*_∞_(0, *z*) = 1 for each *z* ∈ ℝ, and we conclude that

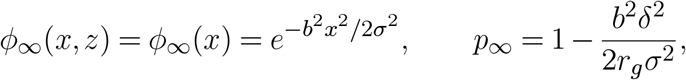

as claimed.

### Evolutionary timescale: Proof that *v* **given by** (6) **satisfies** (1)

We now work in the (slow) evolutionary timescale, and we assume that what happens in the ecological timescale is kept constant to the equilibrium, i.e. we replace *ϕ*(*t, x, z*) with *ϕ*_∞_(*x*−*z*/*b*), namely:

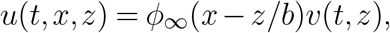

where *u* satisfies the gradient equation (3). We now compute:

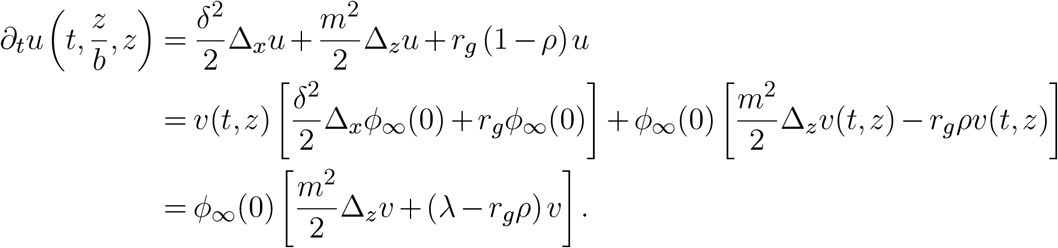

We recall from the previous paragraph, that 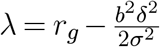 is the principal eigenvalue associated with the operator *L*. Dividing both sides by *ϕ*_∞_(0) gives:

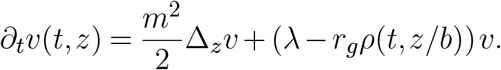

Now, letting 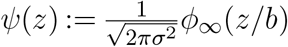 and performing a change of variable, we have *∫*_ℝ_ *ψ* = 1, and:

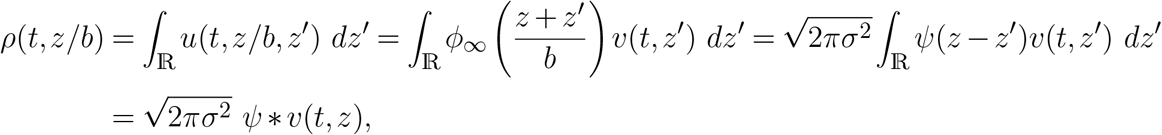

so *v* solves the equation of (7), as claimed.

## C Computations for Section 5.4

### Proof of Equation (10)

Let us write **V**_*t*_ = **V**^*eq*^ + **h**_*t*_ for small **h**_*t*_, with 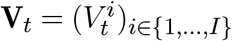 and 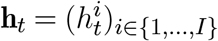. We then have, since 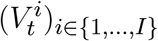 solves (9):

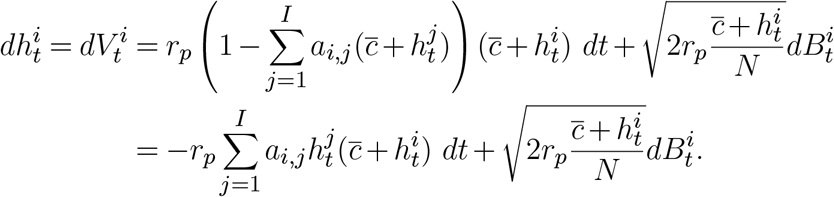

Assuming that **h** is small, this can be approximated, in the first order, by:

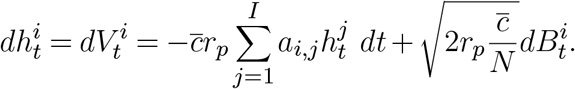

This corresponds exactly to (10).

#### Proof of Equation (13)

We start from the condition *λ*[*η*]*N* = *C* and we prove (13). Using the expression of the limit of *λ*[*η*], the above condition becomes:

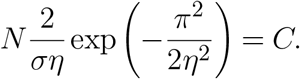

Dividing both sides by 2*N*/*σπ* and taking minus the square gives:

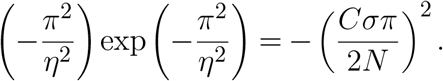

Now, recall that *η* ∈ (0, *π*), so 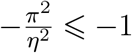, and *N* is sufficiently large so that 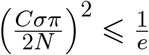. A standard analysis shows that for each *a* ∈ (0, 1/*e*], the equation *xe*^*x*^ = −*a* has exactly one solution below −1. This solution is denoted by *x* = *W*_−1_(*a*) (the function *W*_−1_ is called the *branch of index* −1 *of the Lambert W function*). We deduce from the above equation that

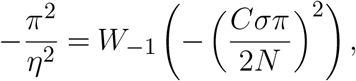

which leads to (13).

#### Proof of Equation (15)

The asymptotic expansion of *W*_−1_ is, for *z* ∈ [−1/*e*, 0) close to 0:

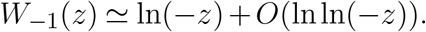

As a consequence, for large *N*, we can make the approximation:

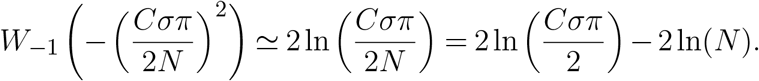

From the above, we deduce

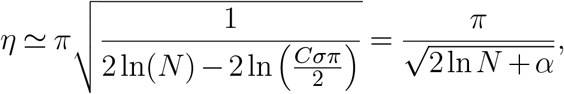

with 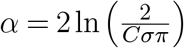.

## References

Alfaro, M., Berestycki, H. & Raoul, G. (2017) The effect of climate shift on a species submitted to dispersion, evolution, growth, and nonlocal competition. SIAM Journal on Mathematical Analysis 49, 562–596 doi: 10.1137/16M1075934.

Alfaro, M., Coville, J. & Raoul, G. (2013) Travelling waves in a nonlocal reaction-diffusion equation as a model for a population structured by a space variable and a phenotypic trait. Communications in Partial Differential Equations 38, 2126–2154 doi: 10.1080/03605302.2013.828069.

Barabás, G. & Meszéna, G. (2009) When the exception becomes the rule: The disappearance of limiting similarity in the Lotka–Volterra model. Journal of Theoretical Biology 258, 89–94 doi: 10.1016/j.jtbi.2008.12.033.

Barton, N.H. (1999) Clines in polygenic traits. Genetics Research 74, 223–236 doi: 10.1017/S001667239900422X.

Berestycki, H., Nadin, G., Perthame, B. & Ryzhik, L. (2009) The non-local Fisher-KPP equation: travelling waves and steady states. Nonlinearity 22, 2813–2844 doi: 10.1088/0951-7715/22/12/002.

Boutillon, N. (2025) Qualitative properties of the spreading speed of a population structured in space and in phenotype. Journal de Mathématiques Pures et Appliquées. Neuvième Série 204, Paper No. 103804, 38 doi: 10.1016/j.matpur.2025.103804.

Boutillon, N. & Rossi, L. (2025) Reaction–diffusion model for a population structured in phenotype and space: I. criterion for persistence. Nonlinearity 38, 045019 doi: 10.1088/1361-6544/adbda4.

Britton, N.F. (1989) Aggregation and the competitive exclusion principle. Journal of Theoretical Biology 136, 57–66 doi: 10.1016/S0022-5193(89)80189-4.

Champagnat, N., Ferrière, R. & Méléard, S. (2006) Unifying evolutionary dynamics: from individual stochastic processes to macroscopic models. Theoretical population biology 69 doi: 10.1016/j.tpb.2005.10.004.

Champagnat, N. & Méléard, S. (2007) Invasion and adaptive evolution for individual-based spatially structured populations. Journal of Mathematical Biology 55, 147–188 doi: 10.1007/s00285-007-0072-z.

Dieckmann, U. & Doebeli, M. (1999) On the origin of species by sympatric speciation. Nature 400, 354–357 doi: 10.1038/22521.

Doebeli, M. & Dieckmann, U. (2003) Speciation along environmental gradients. Nature 421, 259–264 doi: 10.1038/nature01274.

Fang, J. & Zhao, X.Q. (2011) Monotone wavefronts of the nonlocal Fisher-KPP equation. Nonlinearity 24, 3043–3054 doi: 10.1088/0951-7715/24/11/002.

Felsenstein, J. (1977) Multivariate normal genetic models with a finite number of loci. Proceedings of the International Conference on Quantitative Genetics. p. 227–245 Iowa State Univ. Press, Ames, IA.

Fouqueau, L. & Roze, D. (2021) The evolution of sex along an environmental gradient. Evolution 75, 1334–1347 doi: 10.1111/evo.14237.

Genieys, S., Volpert, V. & Auger, P. (2006) Pattern and waves for a model in population dynamics with nonlocal consumption of resources. Mathematical Modelling of Natural Phenomena 1, 65–82 doi: 10.1051/mmnp:2006004.

Haldane, J.B.S. (1919) The combination of linkage values, and the calculation of distance between the loci of linked factors. Journal of Genetics 8, 299–309.

Haldane, J.B.S. (1948) The theory of a cline. Journal of Genetics 48, 277–284 doi: 10.1007/BF02986626.

Hamel, F. & Ryzhik, L. (2014) On the nonlocal Fisher-KPP equation: steady states, spreading speed and global bounds. Nonlinearity 27, 2735–2753 doi: 10.1088/0951-7715/27/11/2735.

Hernández-García, E., López, C., Pigolotti, S. & Andersen, K.H. (2009) Species competition: coexistence, exclusion and clustering. Philosophical transactions. Series A, Mathematical, physical, and engineering sciences 367, 3183 doi: 10.1098/rsta.2009.0086.

Kirkpatrick, M. & Barton, N.H. (1997) Evolution of a species’ range. The American Naturalist 150, 1–23 doi: 10.1086/286054.

Kulmuni, J., Butlin, R.K., Lucek, K., Savolainen, V. & Westram, A.M. (2020) Towards the completion of speciation: the evolution of reproductive isolation beyond the first barriers. Philosophical Transactions of the Royal Society B: Biological Sciences 375 doi: 10.1098/rstb.2019.0528.

Leimar, O., Doebeli, M. & Dieckmann, U. (2008) Evolution of phenotypic clusters through competition and local adaptation along an environmental gradient. Evolution 62, 807–822 doi: 10.1111/j.1558-5646.2008.00334.x.

Leimar, O., Sasaki, A., Doebeli, M. & Dieckmann, U. (2013) Limiting similarity, species packing, and the shape of competition kernels. Journal of Theoretical Biology 339, 3–13 doi: 10.1016/j.jtbi.2013.08.005.

May, R.M. (1969) Species packing, and what competition minimizes. Proceedings of the National Academy of Sciences 64, 1369–1371 doi: 10.1073/pnas.64.4.1369.

May, R.M. & MacArthur, R.H. (1972) Niche overlap as a function of environmental variability. Proceedings of the National Academy of Sciences 69, 1109–1113 doi: 10.1073/pnas.69.5.1109.

Nadin, G., Rossi, L., Ryzhik, L. & Perthame, B. (2013) Wave-like solutions for nonlocal reaction-diffusion equations: a toy model. Mathematical Modelling of Natural Phenomena 8, 33–41 doi: 10.1051/mmnp/20138304.

Pigolotti, S., López, C., Hernández-García, E. & Andersen, K.H. (2010) How Gaussian competition leads to lumpy or uniform species distributions. Theoretical Ecology 3, 89–96 doi: 10.1007/s12080-009-0056-2.

Polechová, J. & Barton, N.H. (2005) Speciation through competition: a critical review. Evolution 59, 1194–1210 doi: 10.1111/j.0014-3820.2005.tb01771.x.

Polechová, J. & Barton, N.H. (2015) Limits to adaptation along environmental gradients. Proceedings of the National Academy of Sciences 112, 6401–6406 doi: 10.1073/pnas.1421515112.

Prevost, C. (2004) Applications des équations aux dérivés partielles aux problèmes de dynamique des populations et traitement numérique. Phd thesis Université d’Orléans.

Rogers, T., McKane, A.J. & Rossberg, A.G. (2012) Demographic noise can lead to the spontaneous formation of species. Europhysics Letters 97, 40008 doi: 10.1209/0295-5075/97/40008.

Roze, D. (2014) Selection for sex in finite populations. Journal of Evolutionary Biology 27, 1304–1322 doi: 10.1111/jeb.12344.

Sasaki, A. (1997) Clumped distribution by neighbourhood competition. Journal of Theoretical Biology 186, 415–430 doi:10.1006/jtbi.1996.0370.

